# Potassium induces conformational changes in the Sabiá virus spike complex

**DOI:** 10.1101/2024.08.22.609195

**Authors:** Hadas Cohen-Dvashi, Michael Katz, Ron Diskin

## Abstract

Hemorrhagic fever viruses from the *Arenaviridae* are a source of concern due to their potential to cause lethal outbreaks and the lack of effective therapeutics. Several Clade-B viruses, which are endemic to the Americas, are pathogens that sporadically infect humans following zoonotic transmission from small rodents. Brazilian hemorrhagic fever, caused by the Sabiá virus (SBAV), is one of several such diseases from Clade-B arenaviruses. Despite their importance and the risks they impose, many fundamental questions remain regarding their biology and function. Here, we present the structure of the spike complex from the Sabiá virus, which mediates viral attachment and entry to the host cells. Our study reveals two distinct conformational states of the spike, representing its native closed state and an open state that it assumes during cell entry. We show that potassium, in combination with acidic pH, promotes the opening of the spike, which is required for achieving efficient cell entry. This structure further informs us about the architecture of Clade-B arenaviral spikes and how they vary from spikes of other members of the *Arenaviridae*.

## Introduction

Sabiá virus (SBAV), also known as *Mammarenavirus brazilense*, is a human pathogen that causes Brazilian hemorrhagic fever ^1^. The virus was discovered in 1990 and has a fatality rate of ∼50% of symptomatic cases in humans ^2^. It is a ‘Clade B’ mammarenavirus that comprises part of the ‘New World’ (NW) group within the *Arenaviridae*. Besides SBAV, there are several other known disease-causing Clade B mammarenaviruses. These viruses circulate in various small animal hosts, mostly rodents ^3^, and are endemic to South American countries. Following close contact with animals or with their excreta, these viruses can be zoonotically transmitted to humans, resulting in various clinical symptoms ranging from mild fever diseases to severe and often deadly hemorrhagic fevers ^4^. Additional notable disease-causing Clade B mammarenaviruses include the Junín virus (JUNV), the Machupo virus (MACV), the Guanarito virus (GTOV), and the Chapare virus (CHAPV), which are the etiology agents of the Argentine, the Bolivian, the Venezuelan, and the Chapare hemorrhagic fevers, respectively. Documented nosocomial transmission through aerosol of some of these viruses ^5,6^ emphasizes the risk for outbreaks and the potential use of these viruses as biological weapons ^7,8^. Besides supportive care, therapy is limited to ribavirin or convalescent plasma, which have limited efficacy ^9^. Therefore, most of these disease-causing Clade B mammarenaviruses are classified as high-priority pathogens by the National Institute of Allergy and Infectious Diseases (NIAID), calling for basic research and the development of new therapeutics.

Arenaviruses are small, enveloped, single-stranded RNA viruses ^4,10^. They utilize class-I trimeric glycoprotein spike complexes to engage with their cellular receptors and enter their host cells ^11^. In the *Mammarenavirus* genus, these spikes are translated as a single polypeptide chain. Three such chains trimerize, forming the glycoprotein precursor complex (GPC). To assume its mature state, the GPC needs to be processed twice, by a signal peptidase (SPase) and the subtilisin kexin isozyme-1/site-1 (SKI-1) protease ^12-14^, producing three different subunits: a stable signal peptide (SSP), a glycoprotein 1 (GP1) receptor binding domain, and a glycoprotein 2 (GP2) membrane fusion domain (Fig. 1a). All clade B mammarenaviruses use transferrin receptor protein 1 (TfR1) as their cellular receptor ^3,15,16^. Interestingly, isolated GP1 domains from these clade B mammarenaviruses readily bind TfR1 ^17,18^. This ability significantly differs from mammarenaviruses that utilize other receptors. Arenaviruses from the ‘Old World’ (OW) group, like the Lassa virus (LASV) that is endemic to West Africa or the lymphocytic choriomeningitis virus (LCMV) that is globally distributed, use α-dystroglycan as an entry receptor ^19,20^. Rather than directly attaching to α-dystroglycan, these OW mammarenaviruses bind matriglycan, which is a long linear glycan that decorates α-dystroglycan ^21^. The binding site of matriglycan is quaternary and only forms in the context of the intact trimeric spike, as it spans more than a single GP1 domain ^22^. The Lujo virus (LUJV), which is a distinct mammarenavirus, that does not belong to the classical NW or OW groups of mammarenaviruses ^23^, utilizes neuropilin 2 (NRP2) as a cellular receptor ^24^. While isolated GP1 domains of LUJV retain some binding to NRP2 ^25^, the complete binding site is also quaternary, forming by the association of neighboring GP1 domains in the context of the spike ^26^. Furthermore, the cell entry of LASV, LCMV, LUJV, and perhaps other OW mammarenaviruses involves pH-induced dissociation from their respective primary cell entry receptor and the engagement with a secondary, intracellular receptor ^24,27,28^, which serves as a triggering factor for membrane fusion ^29^. Such pH-induced receptor release has not been reported so far for TfR1-tropic mammarenaviruses, nor has the use of secondary intracellular receptors. Taken together, the functional diversity suggests substantial differentiation of arenaviral spike complexes.

**Fig. 1.**
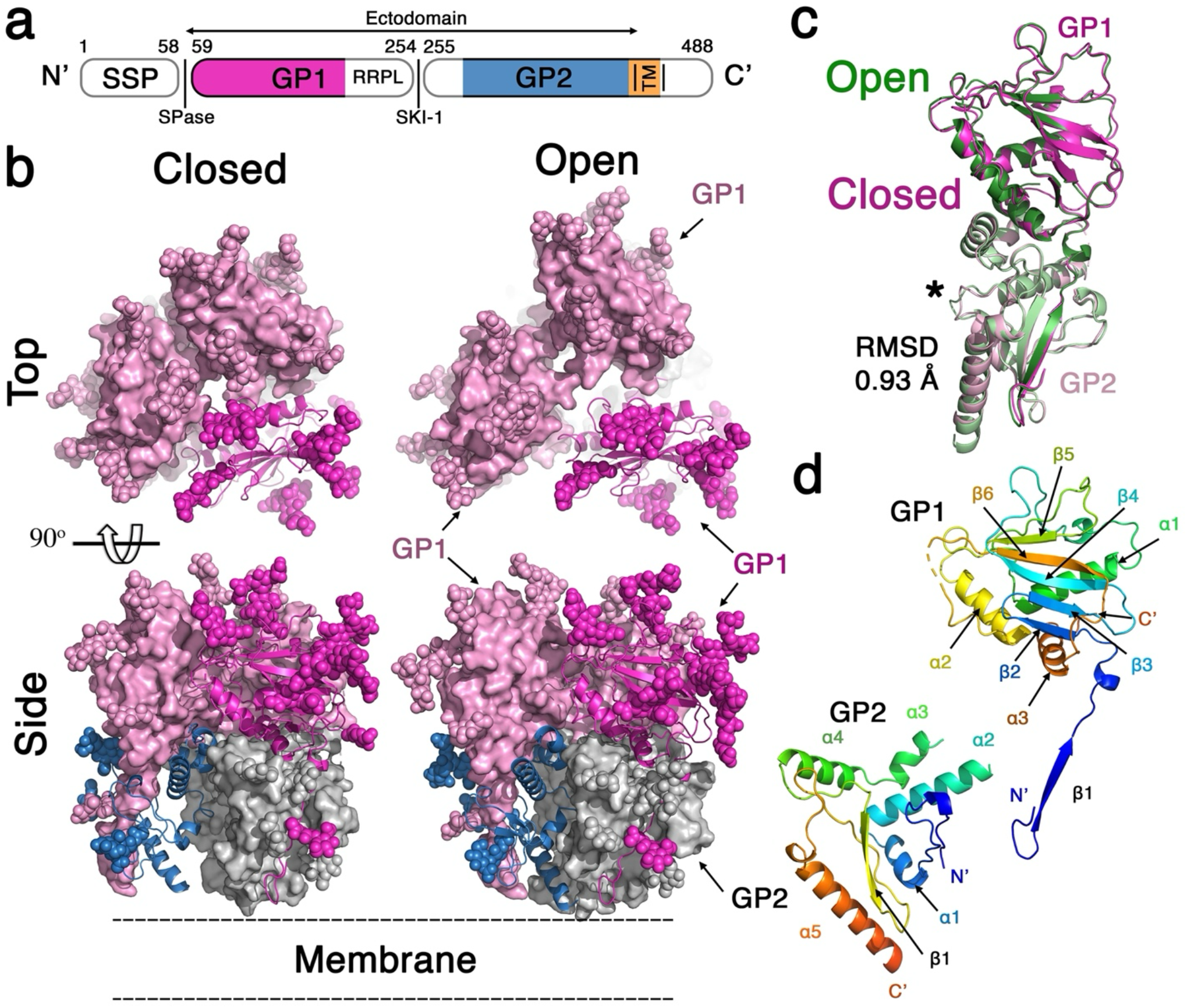
The overall structure of the SBAV spike complex. **a**. Schematic diagram (not to scale) showing the organization of the GPC protomer. Colored regions illustrate the regions that were modeled in the structure. Part of the TM region (orange) was resolved in the structure of the H157M mutant (see below) **b**. The SBAV spike adopts two distinct conformations. The open and closed conformations of the spike are shown using ‘top’ and ‘side’ views. Pink and grey surface representations show the GP1 and GP2 domains, respectively. One GP1 (purple) and one GP2 (blue) are highlighted using ribbon representation. N-linked glycans are indicated with spheres, and the orientation of the spike with respect to the membrane is indicated. **c**. Superimposition of GP1/GP2 protomers from the open (green) and closed (pink) conformations. RMSD based on 316 shared Cα atoms is indicated. An asterisk indicates a GP2 loop (*i*.*e*., α1-α2 loop) that assumes different conformations in the two states. **d**. Ribbon representation of the GP1 and GP2 domains, rainbow-colored from the N’ (blue) to the C’ (red). The termini and the secondary structure elements are indicated.

While structural data for the antigenic appearance of the LASV and LCMV spikes ^30-34^ and for the native structures of LASV and LUJV spike complexes ^22,26^ are available, we currently lack such information for spikes of NW mammarenaviruses. The available information is limited for isolated GP1 domains ^35-37^ or for a GP2 domain in its post-fusion state ^38^. Critical questions remain regarding the organization, appearance, and function of the TfR1-tropic mammarenavirus spike complexes. Here, we present a cryo-EM investigation of the spike complex of the NW mammarenavirus SBAV as a representative Clade B mammarenavirus. This study elucidates the spike’s structure, two distinct functional states that it adopts, and the role of a metal ion in its function.

## Results

### Two distinct prefusion states of the spike

Building on our previous success in producing fulllength spike complexes of LASV and LUJV for structural studies ^22,26^, we used a similar approach to isolate the complete spike complex of the SBAV by transiently expressing the SBAV GPCs fused to a Flag-tag at their C-terminus in HEK293F cells and then solubilizing the cellular membranes in detergent. Single particle cryo-EM analysis of the protein sample revealed the presence of two distinct threefold-symmetric states (Extended Data Fig. 1). The first state, which represents an “open” conformation of the spike, provided a 2.9 Å resolution map, and the second state, which shows a “closed” conformation was resolved to 2.6 Å resolution (Extended Data Fig. 1). For both states, the quality of the maps was good (Extended Data Fig. 2a), allowing us to unambiguously model the entire ectodomains that consist of three GP1 and three GP2 subunits, each (Fig. 1b, Extended Table 1). We did not model the TM regions in either case. Low-pass filtering of the EM maps revealed some density for the TM region in the closed but not in the open state (Extended Data Fig. 2b). However, this density was not sufficiently detailed to allow modeling. Nevertheless, the TM region was subsequently partially resolved in a map of a mutated spike (see below).

Looking at the vertical profile of the spike, we can see that the apical region, which consists of the three GP1 domains, is wider, and the GP1s are farther apart from one another in the open state compared to the closed state (Fig. 1b). A ‘top’ view of the spike reveals that in the open state, there are substantial gaps forming between the three GP1 domains, which are not present in the closed state (Fig. 1b). Interestingly, GP1/GP2 pairs from the closed and open states are almost identical, with a root mean square deviation (RMSD) value of 0.93 Å for superimposing them on each other (Fig. 1c). This observation indicates that each GP1/GP2 pair is moving as one rigid body in the transition between these states. Local conformational changes between the states are scarce. The most noticeable conformational change is in a loop (Fig. 1c) that connects α1 with α2 of GP2 (Fig. 1d). Otherwise, the structures of both GP1 and GP2 domains and the relative orientation between the domains within each pair are maintained.

The GP1 receptor binding domain starts with a long β-strand (β1) at its N-terminus (Fig. 1d, Extended data Fig. 3a) that pairs with the β1 strand of GP2, in a form that likely stabilizes the GP2 domain (Extended Data Fig. 3b). The GP1 has a central β-sheet flanked by helices and loops (Fig. 1d). The GP1 of SBAV adopts the same fold and a similar structure to solved GP1 domains of other NW mammarenaviruses (Extended Data Fig. 4). Compared with the GP1 domains of JUNV ^37^, WWAV ^35^, and MACV ^36^, the latter is closest in structure to SBAV (Extended Data Fig. 4). Besides the β1 strand, the GP2 domain only consists of α-helices and loops (Fig. 1d). Comparing the fold of the GP2 domain to the previously determined structures of the LUJV and LASV spike complexes ^22,26^ reveals similar organization (Extended Data Fig. 5a). The density for the first 17 residues of GP2 that contain the fusion peptide was missing, but the following residues that make the fusion loop were modeled (Extended Data Fig. 5b). In the structure of the LUJV spike ^26^, the N-terminal region that contains the fusion peptide was also only partially resolved. In contrast, in the structure of the LASV spike, the entire N-terminus of GP2 was visible, and it extends into the spike’s core, reaching the spike’s 3-fold symmetry axis ^22^.

### The closed vs. the open states

In the closed state, the three GP1 domains closely associate with each other. Each GP1 domain has two distinct interfaces with the neighboring GP1 domains at the clockwise and counterclockwise locations (Fig. 1b). The buried surface area (BSA) at each such interface is ∼1205 Å^2^ (587 Å^2^ on the one monomer and 618 Å ^2^ on the other), reaching a total of ∼3615 Å^2^ BSA for the association of the GP1 domains in the trimer. These interfaces between the GP1 domains are very polar (Fig. 2a). Each GP1 has two patches that have opposite and complementary surface electrostatic potentials (Fig. 2a). The positive patch mostly forms on α1 helix of GP1, and the negative patch involves two loops: α1/β5 and β5/α2 (Fig. 1d and 2a). Key interactions include salt bridges between the Arg246/Asp158 and Arg149/Asp154 pairs, burial of the His157 side chains into a fairly hydrophobic pocket (Fig. 2a), and a stacking cation-π interaction between His186 and Arg139 (Extended Data Fig. 6). In addition to these interactions, several hydrogen bonds involving both side chain and main chain groups further stabilize the interaction (Fig. 2a).

**Fig. 2.**
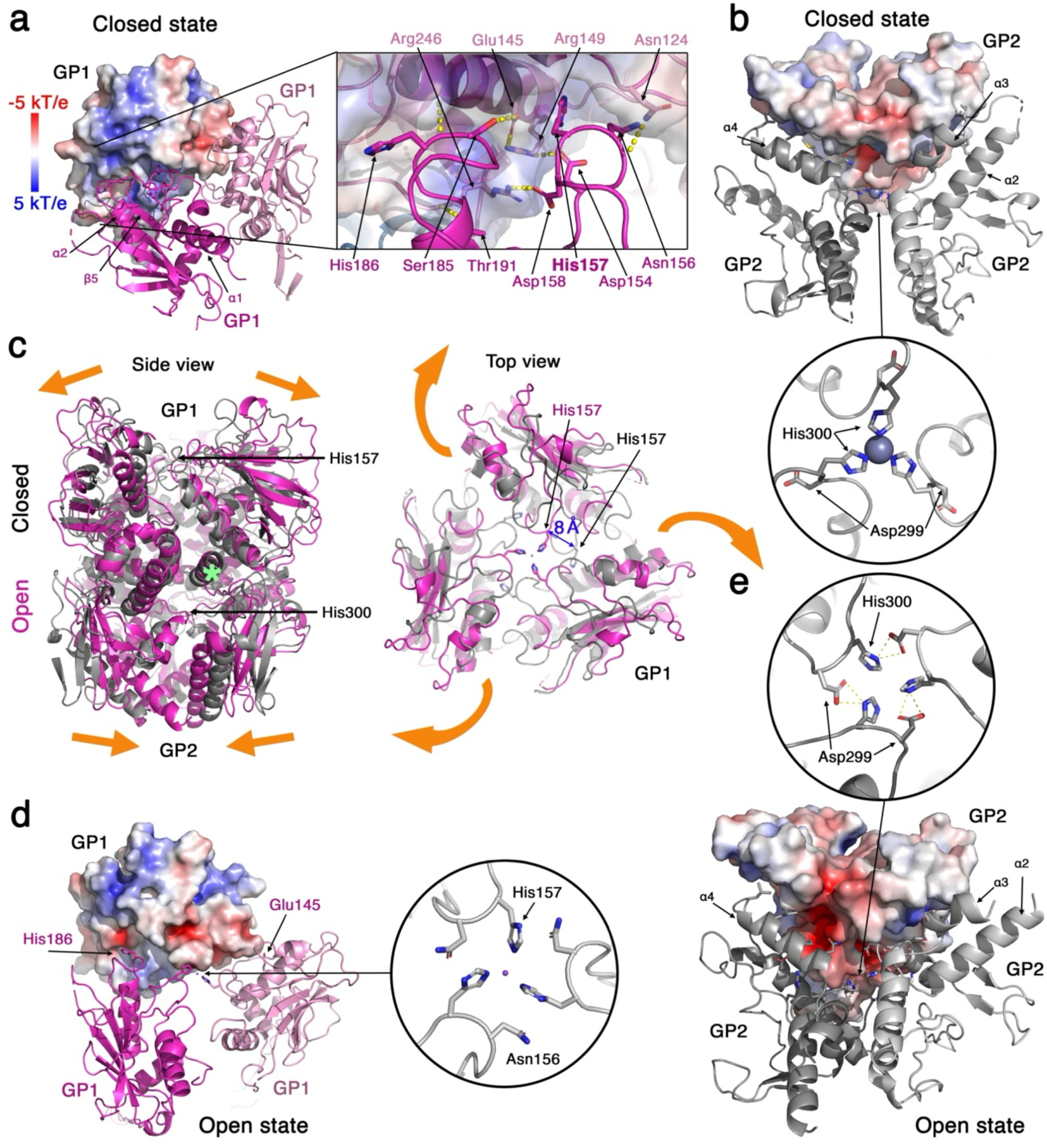
The open and closed states of the spike. **a**. The GP1/GP1 interactions at the closed state. One GP1 subunit is shown with a surface representation colored by the local electrostatic potential (blue 5 kT/e, red -5 kT/e). The inset highlights some of the polar interactions between the GP1 domains. **b**. The GP2/GP2 interfaces in the closed state. One GP2 subunit is shown as a surface colored with local electrostatic potential. The inset shows the metal coordination site that is formed by three His300 residues at the spike’s three-fold symmetry axis. **c**. The transition from closed to open states. The SBAV spike in the closed (grey) and open (purple) states is shown using side and top views. Orange arrows illustrate the movement of the GP1/GP2 pairs. The green asterisk shows the pivot point on one of the GP1/GP2 pairs. In the top view, His157 is shown as sticks, and the extant in which its Cα atom moves is indicated. The position of His157 in GP1 and His300 in GP2 are indicated in the side view. **d**. The GP1/GP1 interface of the open state. Key interacting residues and the electrostatic potential of one GP1 subunit are shown. The inset shows a close-up view of the metal binding site formed by His157. **e**. The GP2/GP2 interfaces in the open state. One GP2 subunit is shown as a surface colored with local electrostatic potential. The inset shows the reformed structure by His300 and Asp299 at the spike’s three-fold symmetry axis.

Analyzing the interaction between the GP2 subunits in the closed state is limited to the modeled region that is missing the membrane-proximal region of the α5 helices, as well as the transmembrane helices (Fig. 1b Extended Data Fig. 1). An improved reconstruction for this region in the closed state was obtained for a H157M mutant of the spike (see details below), revealing the complete α5 helices and part of the transmembrane helices (Extended Data Fig. 7 & Extended Data Table 1). The interaction between the GP2 domains is dominated by hydrophobic interactions (Fig. 2b). Each GP2/GP2 interface includes ∼1412 Å^2^ of BSA (682 Å^2^ on the one monomer and 730 Å^2^ on the other) with a total of 4236 Å^2^ for the entire spike. A central part of the interface includes the α4 helix on one GP2 subunit that interacts with a α3 helix on the second GP2 subunit (Fig. 2b). The α5 helices from different GP2 subunits form mutual interactions involving Tyr414 on one helix that intercalates between Gln418 and Leu423 and stabilizes a 90° turn on the other helix. Tyr414 further forms a hydrogen bond with the main-chain carbonyl of Pro422 that serves as a helix-breaker (Extended Data Fig. 8). Another important interaction between the GP2 domains in the closed state is a metal coordination site (Fig. 2b). The coordination site forms at the spike’s three-fold symmetry axis by three His300 residues that are located on a loop connecting α1 with α2 helices. The exact identity of this metal ion is unknown. We nevertheless modeled a zinc ion in this site based on its availability in the buffer and the reasonable score that it gets from the CheckMyMetal server ^39^.

Superimposing the open and closed states of the SBAV spike complex illustrates the relative movements of the GP1/GP2 pairs during the transition between the states (Fig. 2c). Each GP1/GP2 pair performs a complex movement around a central pivot point. This motion includes a movement of the GP1 domains away from each other, a movement of the lower part of the GP2 domains toward each other, and a clockwise rotation of each GP1/GP2 pair as viewed from the top (Fig. 2c). The transition is illustrated by morphing between the two states, as can be seen from a side view (Extended Data Movie 1) and from a top view (Extended Data Movie 2). The interaction of the GP1 domains in the open state has a substantially reduced BSA compared with the closed state. At each GP1/GP1 interface, there is ∼518 Å^2^ of BSA (257 Å ^2^ on the one monomer and 261 Å ^2^ on the other), with a total of ∼1554 Å^2^ for all the GP1 interfaces. Two key interactions contribute to the GP1 interfaces in the open state: His157, which interacts within a hydrophobic pocket in the closed state (Fig. 2a), moves 8 Å away (Fig. 2c) and creates a new metal coordination site at the spike’s threefold symmetry axis (Fig. 2d). Asn156 contributes to this interaction by forming van Der Waals interactions with the neighboring His157 (Fig. 2d). The second key interaction is formed by His186 that breaks from its cation-π interaction with Arg139 in the closed state (Extended Data Fig. 6) and form a new interaction with a negatively-charge region on the neighboring GP1 subunit (Fig. 2d) made by Glu145 (Extended Data Fig. 9). Interestingly, the transition between the closed and open states likely requires the protonation of His186 since the interaction with Arg139 can only occur when His186 is not protonated, and the interaction with Glu145 will be favored when His186 is protonated.

In contrast to the marked reduction in BSA between the GP1 subunits upon spike opening, the BSA at the interfaces between the GP2 subunits is not changing substantially. Each GP2/GP2 interface in the open state spans ∼1392 Å^2^ (675 Å ^2^ on one subunit and 717 Å ^2^ on the other), with a total of ∼4176 Å ^2^ for all three interfaces (with the caveat that the TM helices are no longer visible in this state, and their contribution is not calculated). However, the actual interactions that the GP2 subunits form do change signiQicantly. The α5 helices, which are curved in the closed state (Extended Data Fig. 8), become straight and approach each other in the open state (Fig. 2c & Extended Data Fig. 10a). A salt-bridge interaction form between Lys412 and Asp403 (Extended Data Fig. 10a), as well as hydrophobic interactions between Ile415 on one helix and Leu406 and Leu410 on the other (Extended Data Fig. 10b). The α4 helix makes additional contacts with the neighboring subunit that include polar interactions (Fig. 2e). Most noticeable, the α1/α2 loop of GP2 adopts a different conformation in the open state (Fig. 1c & Fig. 2e). In the open state, His300 no longer coordinates a metal ion like in the closed state (Fig. 2b), and now forms polar interactions with Asp299 that completely alter the orientation of its side chain (Fig. 2b, Fig. 2e). Similarly to the interaction of His186 with Glu145 (Extended Data Fig. 9), the interaction of Asp299 with His300 likely requires the protonation of the latter.

### The functional significance of the open state

Our structural analysis indicates that both His186 and His300 are likely protonated in the open state of the spike, suggesting a connection between the spike opening and acidic conditions. Interestingly, the SBAV spike sample was produced in a pH 8.0 buffer solution. Hence, while acidic pH likely favors the open state, additional factors contribute to the opening of the spike, even in alkaline conditions. Metal coordination by His157 in the open state (Fig. 2d) may be one such factor. Multiple sequence alignment of various GPCs from NW mammarenaviruses reveals that position 157 in SBAV is not conserved (Extended Data Fig. 11). MACV, which has the most similar GP1 to SBAV (Extended Data Fig. 4), has a methionine residue instead of a histidine in position 157 (Extended Data Fig. 11). Methionine in this position is predicted to fit inside the hydrophobic pocket in the closed state of the spike (Extended Data Fig. 12) but cannot coordinate a metal ion in the open state like a histidine, providing a potential means to assess the contribution of metal coordination to the opening of the spike.

Introducing a methionine residue instead of His157 results in a spike that undergoes normal cellular processing similarly to the WT spike (Fig. 3a). Arenacept, which is a TfR1-based immunoadhesin that targets the receptor binding sites of NW mammarenaviruses ^18^, recognizes the H157M mutant on cells at similar levels to the WT spike (Fig. 3b), indicating proper folding and proper presentation of the mutated spike. Surprisingly, cell entry of MLV-based pseudotyped viruses is significantly reduced when the H157M spike is mediating cell entry, compared to the WT (Fig. 3c). Single particle cryo-EM analysis of the H157M mutant confirmed the presence of the spike’s closed state (Extended Data Fig. 13). However, although using the open state’s map of the WT spike (Extended Data Fig. 1) as a target volume for 3D classification of the EM data, we could not obtain any meaningful representation of the open state with the H157M data (Extended Data Fig. 13). Hence, residue 157 in the spike complex of SBAV needs to be able to coordinate a metal ion in order to achieve a stable open state of the spike, which is important for cell entry.

**Fig. 3.**
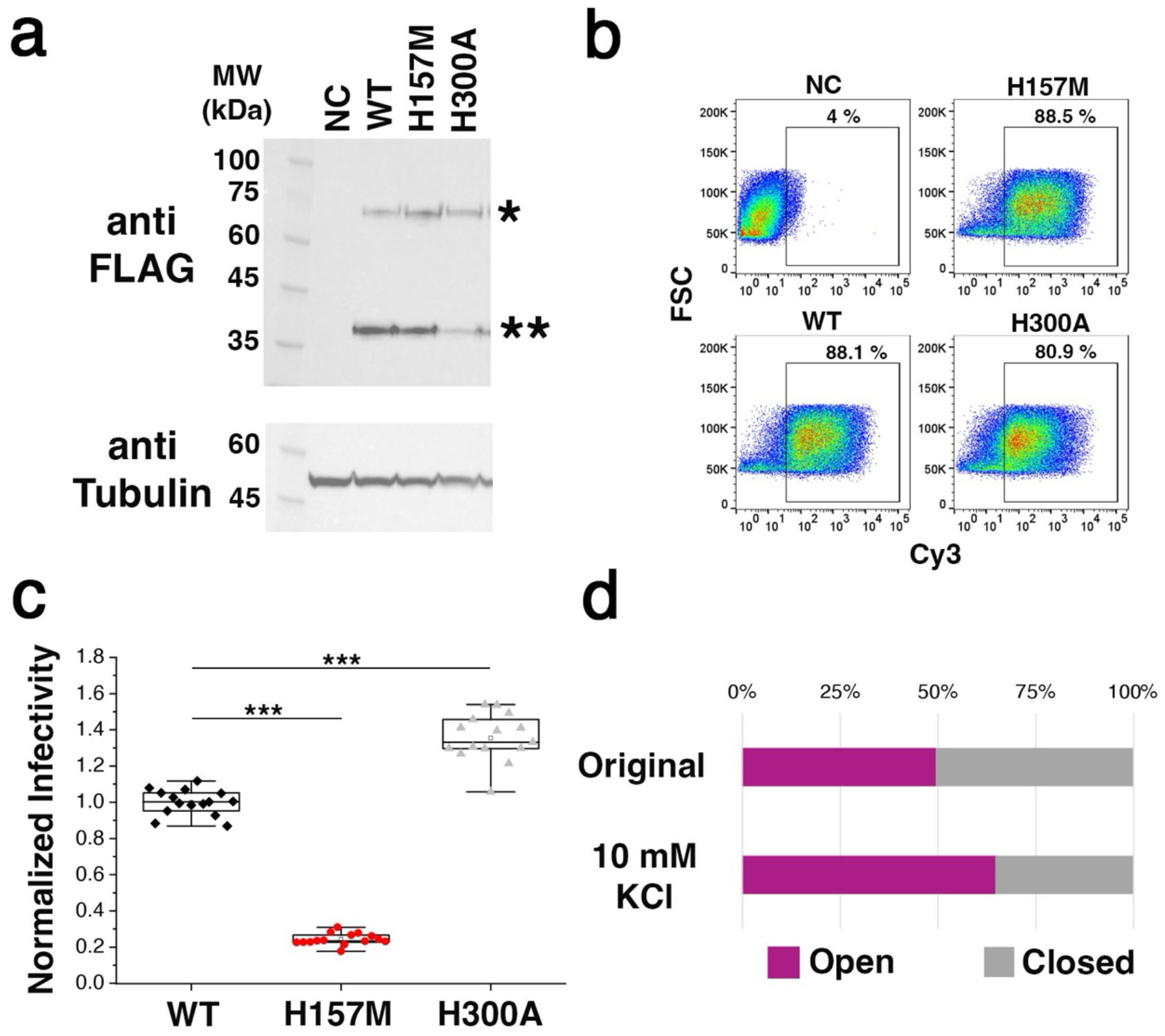
Opening of the spike promotes cell entry. **a**. Western blot analysis of mutated spikes in cell lysates. Negative control (NC) was untransfected cells. ‘*’ and ‘**’ indicate the full GPCs and the processed GP2s, respectively. Proteins were detected with an anti-Flag antibody. Tubulin was used as a load control. **b**. FACS analysis using Arenacept. Cells expressing the indicated spikes or untransfected cells that were used as a negative control (NC) were stained using Arenacept. The percentage of positively stained cells inside the gated area is indicated for each graph. **c**. Infectivity of MLV-based pseudoviruses with mutated spikes. Normalized infectivity levels are indicated. Dots represent technical repeats (n=15). This is a representative experiment out of three independent repeats. Boxes indicate interquartile range; lines indicate median value, and whiskers represent standard deviations. The significance value was calculated using a two-tailed Student’s T-test (‘***’ indicates p<1.0E-9). **d**. The particle distribution between the open (purple) and closed (grey) states for the reconstruction of the original SBAV sample and for the reconstruction after supplementing the sample with 10 mM of potassium.

Single-particle cryo-EM analysis does not provide direct information regarding the identity of elements that contribute to the EM density. Therefore, we do not know which metal ion is coordinated by His157 in the open state (fig. 2d). The potential link of the open state with acidic conditions (Fig. 2e, Extended Data Fig. 9), together with the observed impairment of cell entry (Fig. 3c), when assuming the open state is prevented (Extended Data Fig. 13), indicates that the open state has a functional role during cell entry of SBAV. In the endocytic pathways, besides gradual acidification, the ion composition changes with a most noticeable rise in the concentration of potassium ^40^. Therefore, we hypothesized that His157 coordinates a potassium ion in the open state. To probe this possibility, we supplemented an aliquot of the SBAV spike sample from the same purification batch that was used to determine the structure (Extended Data Fig. S1) with 10 mM of KCl and imaged it using EM. Similar to the original sample, we could readily reconstruct both the open and closed states of the spike (Extended Data Fig. 14). However, unlike the original sample, which had an almost identical number of particles in both states (Fig. 3d & Extended Data Fig. S1), after supplementing the spike with 10 mM of KCl, there were twice the number of particles that belonged to the open state compared to the closed state (Fig. 3d & Extended Data Fig. 14). Hence, potassium shifts the equilibrium toward the open state. While this observation supports the idea that His157 coordinates potassium, it may be that potassium favors the open state by some His157-independent mechanism. To evaluate this possibility, we supplemented the H157M spike with 10 mM potassium and performed single-particle cryo-EM analysis (Extended Data Fig. 7). In the absence of a histidine at position 157, the addition of 10 mM potassium did not result in any interpretable open state (Extended Data Fig. 7). The reconstruction quality of the closed state in these conditions was even better than the original reconstruction of the WT spike (Extended Data Fig. 1), and the excess of particles allowed us to find a subclass of particles that partially revealed the organization of the spike inside the membrane (Extended Data Fig. 7 & Extended Data Table 1). Hence, the presence of potassium promotes the opening of the spike in a mechanism that requires His157.

The coordination of a metal ion by His300 in the closed state (Fig. 2b) may also play an important role in regulating the opening of the spike, as it likely antagonizes the spike’s opening by stabilizing the closed state. His300 is a conserved residue among NW mammarenaviruses (Extended Data Fig. 11), but this is true for the majority of GP2 residues. Introducing an alanine mutation in position 300 of the SBAV GPC results in a spike that manages to express and mature, albeit at a slightly reduced efficiency compared with the WT spike (Fig. 3a). This H300A spike reaches the cell surface and is recognized by Arenacept (Fig. 3b), indicating proper folding and presentation. Interestingly, MLV-based pseudotyped viruses seem to enter cells more efficiently with the H300A spike compared with the WT SBAV spike (Fig. 3c). The mere change in cell entry efficiency establishes a role for this metal coordination site during cell entry. Hence, given that the metal coordination by His300 shifts the equilibrium toward the closed state by stabilizing it, the observed increase in cell entry levels of the H300A mutant indicates that assuming the open state is indeed a critical functional step for the cell entry of SBAV.

### Receptor recognition by the spike complex

The overall structural similarity of the GP1 receptor binding domains among NW mammarenaviruses (Extended Data Fig. 4) allows us to utilize a previously determined structure of the soluble MACV GP1 bound to its human TfR1 receptor ^17^ to deduce how TfR1 is recognized in the context of a complete spike by SBAV. Superimposing the MACV GP1/TfR1 model on the SBAV’s spike structure reveals that the TfR1 binding sites on the GP1 domains are oriented away from the threefold symmetry axis (Fig. 4a). This organization allows all three GP1 domains to engage with TfR1 without causing steric interferences. The transition from the closed to the open state changes the relative orientation of the TfR1s but does not create any steric clashes between the TfR1 molecules, indicating that the opening of the spike does not require the release of the receptor. A closer look at the putative SBAV GP1/TfR1 interaction reveals that several N-linked glycans are strategically flanking the entire TfR1 binding site (Fig. 4b). Of the five N-linked glycans near the TfR1 binding site, the glycan attached to Asn88 will have to adopt a different orientation, compared to the one observed in the EM reconstruction, to allow TfR1 binding as it partially blocks the binding site. Interestingly, the Asn88-linked glycan is conserved (Extended Data Fig. 11), and for the clade A/B TfR1-tropic Whitewater Arroyo virus, it was also shown that this glycan has to change orientation in order to allow receptor binding ^35^.

**Fig. 4.**
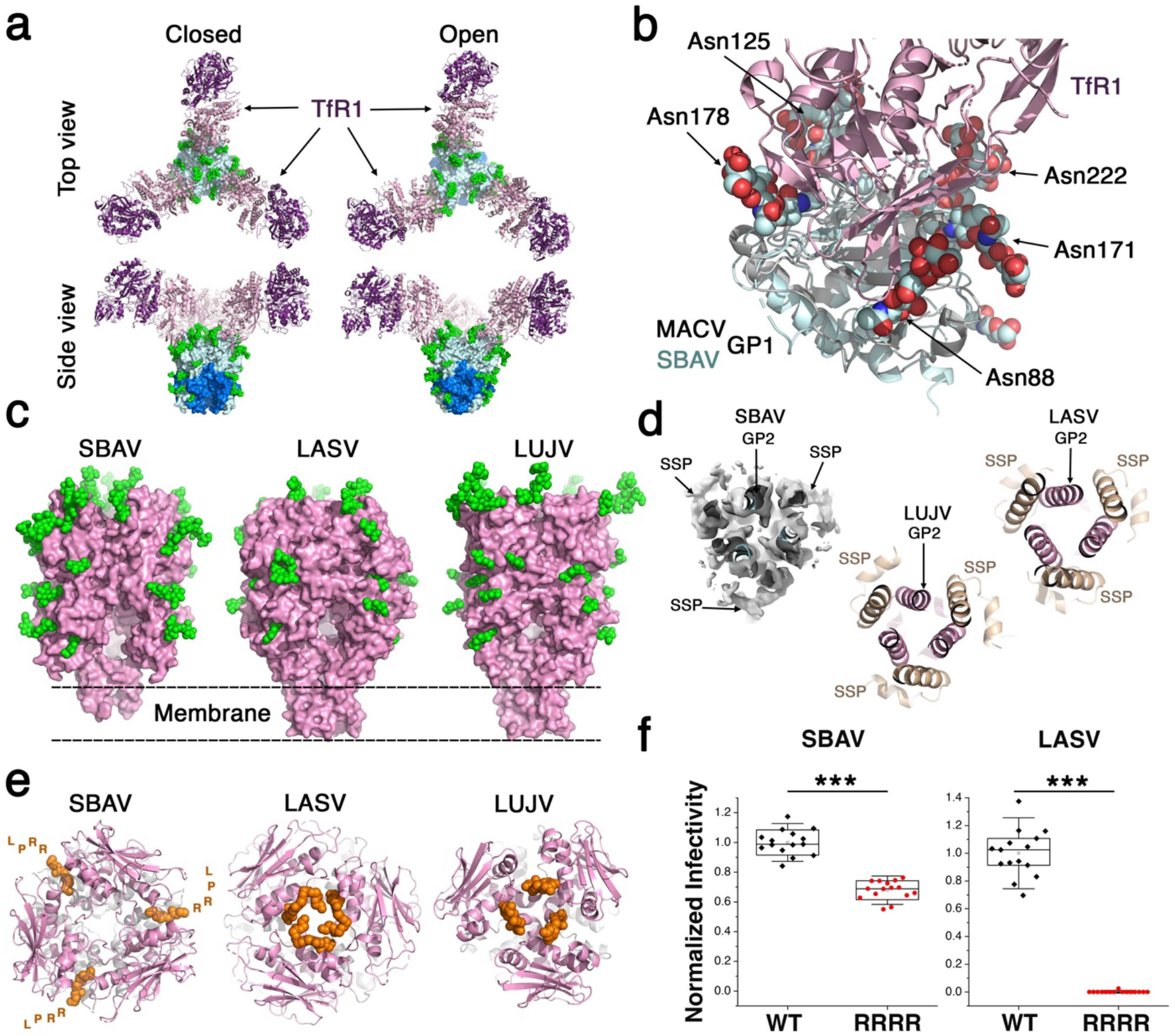
TfR1 recognition by the SBAV spike and the way it compares with other arenaviral spikes. **a**. The SBAV spike can bind three TfR1. A model generated by superimposing the SBAV spike with the MACV GP1/human TfR1 structure (PDB: 3KAS). The model in both the open and closed states is shown using “top” and “side” views in the upper and lower panels, respectively. **b**. A close-up view of the TfR1/SBAV GP1 complex. N-linked glycans are shown as spheres and noted by the residue number to which they are attached. GP1s of SBAV and MACV are shown in cyan and grey ribbon representations, respectively. TfR1 is shown as a pink ribbon. **c**. Overview of the SBAV (H157M, closed state), LASV (PDB: 7PUY), and LUJV (PDB: 8P4T) spike complexes. The spikes are shown using pink surface representations with respect to the rough location of the membrane. N-linked glycans are shown as green spheres. **d**. The transmembrane organization of the spikes. The three spikes are shown from a “bottom” view. The SBAV H157M spike is shown in its EM map (map level=0.035). **e**. Top views of the SBAV, LASV, and LUJV spikes. The C-termini of the GP1s are noted as orange spheres. For SBAV, the last four residues that were not modeled are illustrated by their single-letter codes. **f**. Normalized infectivity of MLV-pseudotyped viruses bearing the indicated spikes. The graphs show representative data from one experiment, which is part of three independent repeats. The dots represent technical (n=15) repeats. Boxes indicate interquartile range; lines indicate median value, and whiskers represent standard deviations. The significance value was calculated using a two-tailed Student’s T-test (‘***’ indicates p<1.0E-11).

### Structural differentiation of mammarenaviral spikes

The current structure of the NW SBAV spike adds to two other arenaviral spikes that were extracted from the membrane, and their entire structures are available, including their transmembrane regions ^22,26^. The GP2 and GP1 domains share the same folds for all the arenaviral proteins that were so far determined. However, the relative organization of the domains and the distribution of N-linked glycans, which closely relates to the receptor usage, are not identical, giving rise to a distinct appearance of the spikes (Fig. 4c). The transmembrane parts of both LASV and LUJV spikes ^22,26^ organize in a similar six-helical bundle configuration, in which the central three helices are of GP2 and the SSP helices incorporate to the bundle as a second layer (Fig. 4d). In the spike of the H157M SBAV, the central three helices are partially visible (Fig. 4c & 4d). Interestingly, in the EM map of the SBAV spike, additional densities that likely belong to the three SSP helices are also partially visible (Fig. 4d). While the density was insufficient to properly model these SSP helices, it does suggest that a six-helical bundle configuration is also adopted by NW mammarenaviruses, implying that this particular organization of the TM helices is a universal feature among mammarenaviruses.

A noticeable difference between the SBAV spike and the spikes of LASV and LUJV is the organization of the GP1s’ C-termini. In both LASV and LUJV spikes, the C-termini of their GP1s, which contain the SKI-1 recognition motifs, are well-ordered through tight interactions with the neighboring GP1 chains (Fig. 4e). These interactions stabilize the trimeric organization of the spikes and are critical for forming the quaternary receptor binding sites of these viruses ^22,26^. In the spike of SBAV, however, the C-termini of the GP1 domains do not make part of the TfR1 binding site and do not interact with neighboring GP1 domains. Instead, the C-termini projects outwards (away from the symmetry axis) from within the GP1/GP1 interface (Fig. 4e). Density is lacking for the last four residues (the “RRPL” SKI-1 recognition motif), indicating structural heterogeneity. Thus, unlike in LASV and LUJV, the GP1 termini in SBAV, following proteolytic cleavage by SKI-I, do not have a structural role *per se* and could, in principle, be adapted to be cleaved by a different cellular protease. Indeed, a mutated SBAV spike in which the “RRPL” SKI-I recognition motif was swapped with an “RRRR”-furin recognition site supports cell entry of MLV-pseudotyped viruses, albeit with some reduced efficiency (Fig. 4f). On the other hand, pseudotyping MLVs with a LASV spike in which the SKI-I recognition motif (which is “RRLL” in the case of LASV) is mutated to an “RRRR” furin site results with non-infectious particles (Fig. 4f). This result is corroborated by the failure of the LASV spike to accumulate in its proteolytic-mature form when the furin site is used (Extended Data Fig. 15), perhaps due to misfolding and clearance of this protein by the ER quality control system. Therefore, the different organization of the arenaviral spikes gives rise to important functional differences.

## Discussion

The *Arenaviridae* is a large and diverse family of viruses, with members that have distinct geographic niches ^10^, show clear genetic diversity ^23^, and use different cell entry strategies ^15,19,24^. The structure of the spike complex from the NW SBAV mammarenavirus reveals, however, features that seem to be preserved among all mammarenaviruses. Compared with the spikes of the genetically distinct LUJV and the OW LASV, the structure of the SBAV spike shows that the overall folds of the SSP and GP2 domains are preserved (Fig. 4d, Extended Data Fig. 5), adding to previous notions that all GP1 receptor binding domains share the same overall fold ^25,35,36,41-43^. All three spikes adopt a six-helical bundle configuration inside the membrane (Fig. 4d), suggesting that the organization of SSP and GP2 inside the membrane is shared by all mammarenaviruses. Despite these similarities, the actual structures of the spikes differ (Fig. 4c & 4e) in a functionally important way. When the C-termini of receptor binding domains are not structurally sequestered following cleavage and hence lack a particular structural role in the mature spike, the sequences may change and alter the cellular protease used for maturation, which in turn could modify viral properties. Acquiring a furin recognition site, for example, may increase virulence. The increase in virulence, as noted in the case of SARS-CoV-2, for example, is enabled by the binding of NRP1, which recognizes exposed furin sites following cleavage and becomes an attachment factor ^44,45^. As revealed in the structure of the SBAV spike, NW mammarenaviruses, in contrast to OW mammarenaviruses, have exposed C-termini and may thus evolve to use different cellular proteases for maturation. The fact that NW mammarenaviruses nevertheless adhere to SKI-1 as their cellular protease ^12^ and that the entry efficiency of furin-cleaved SBAV is apparently reduced (Fig. 4f) suggests other advantages for using SKI-1. The exact cellular compartment where cleavage of various mammarenaviral spikes occurs is fine-tuned by the sequence of the SKI-1 recognition motif ^14^. Thus, cleavage of the SBAV spike by furin upon introducing the “RRRR” motif may occur at a suboptimal cellular location, which somehow hinders proper maturation of the spike.

Interestingly, in the protein sample that we produced, the spike complex of SBAV adopted two distinct conformations, representing closed and open states. Such conformations were not previously observed in the single-particle cryo-EM analyses of LASV ^22^ or LUJV ^26^. However, low-resolution data for the LASV spike did indicate pH-induced conformational changes ^46^, which may be important for engaging with its intracellular receptor ^27,46^. Not only LASV but also LCMV and LUJV need to engage with intracellular receptors to facilitate cell entry ^24,28^, suggesting that their spike complexes may also change conformations during cell entry. The open state of the SBAV spike complex, which can likely be promoted by acidic pH (Fig. 2e, Extended Data Fig. 6, 9) and by coordination of a metal ion with His157 (Extended Data Fig. 13), seems to be a critical step in the cell entry process of SBAV (Fig. 3c). It is not clear why the SBAV spike needs to open for efficient cell entry. Considering the requirement of LASV, LCMV, and LUJV to engage with intracellular receptors for cell entry ^24,27,28^, a compelling idea is that the open state is needed to provide access to a yet-to-be-discovered intracellular receptor that the SBAV is utilizing. Regardless of the functional role of the open state, we hypothesize that in the extracellular environment, the SBAV spike complex will be mostly locked in the closed state to minimize its exposed surfaces that could be targeted by antibodies. Hence, our purification process somehow unleashed the ability of the spike to open. One interesting observation is that unlike in the structures of LASV and LUJV ^22,26^, the transmembrane region of the SBAV spike was mostly disordered (Extended Data Fig. 2b). Taken together with our previous notion that the SSP is a critical stabilizing element of the LASV spike ^22^, it could be that our purification conditions caused the SSP to detach from the spike, which in turn, destabilized the spike and enabled its opening. This speculation will need to be investigated in the future. Regardless, variations in the cell entry mechanisms of mammarenaviruses likely exist, and it is not clear if other viruses can adopt such an open state. His157, for example, promotes spike opening by coordinating a metal ion and likely serves as a pH sensor, but it is not a conserved residue (Extended Data Fig. 11). Therefore, if spikes of other viruses like MACV adopt an open state during cell entry, they may utilize different mechanisms to control opening.

Our data establishes a connection between a functionally important conformation that the SBAV spike complex assumes during cell entry and the presence of potassium. The requirement of potassium for productive cell entry and potassium-induced conformational changes in viral glycoproteins have already been noted before. For the Bunyamwera virus (BUNV), which is a member of the *Orthobunyavirus* genus of the *Peribunyaviridae* family, the presence of potassium in the endosomal pathways is critical for cell entry ^47^. The glycoproteins of the Hazara virus (HAZV), which is a *Nairoviridae* family member, undergo potassium-induced conformational changes when exposed to elevated concentrations of potassium, which also increases its infectivity potential ^48^. An increase in cell entry efficiency following exposure to high potassium concentrations for HAZV is reminiscent of the observed increased infectivity rate for the SBAV H300A mutant (Fig. 3c) that likely shifts to the open state more easily. Recently, a genetic screen identified potassium channels as being important for the cell entry of the OW LCMV mammarenavirus ^49^. It was postulated that in the case of LCMV, the potassium acts inside the virion to modulate the interaction of the matrix protein Z with the viral nucleoprotein and, by that, mediates uncoating ^49^. Considering our results, the dependency of LCMV on the presence of potassium may further result from a structural effect that potassium has on the spike complex. Whether LCMV or other mammarenaviruses also change conformation following the binding of potassium, like in the case of SBAV, is an open question. Interestingly, although belonging to different viral families, SBAV, HAZV, and BUNV are all viruses within the *Bunyavirales* order. It may be that sensing and utilizing potassium during cell entry is an ancient trait of Bunyaviruses that is still utilized by some distinct members of this viral order.

## Materials and Methods

### Cloning and expression vectors

Codon-optimized SBAV GPC (UniProt Seq. H6V7J2) was chemically synthesized (Genescript) and subcloned fused to a C-terminal Flag tag into pcDNA3.1 using BamHI–NotI restriction sites. Histidine mutants (H157M, H300A) and SKI-I to Furin (P253R+L254R) mutant of SBAV-FLAG GPC were generated by site-directed PCR mutagenesis using KAPA HiFi hot-start polymerase (Roche). The generation, expression, and purification of sAD-Neotomis TfR1 Fc (Arenacept) were previously described ^18^, as well as the cloning of LASV-FLAG GPC ^22^.

### SBAV GPC production and purification

Expression of the full-length FLAG-tagged SBAV GPC was carried out in HEK293F cells (Invitrogen) using FreeStyle Medium (Life Technologies). Cells were grown to a density of approximately 1.0 × 10^6^ cells per ml before transfection. HEK293F cells were transfected using 40 kDa polyethylenamine (PEI-MAX) (Polysciences) at 1 mg ml^−1^, pH 7 with DNA at a ratio of 1:3 (DNA:PEI solution). The SBAV GPC-expressing cells were collected at 48 h post-transfection by centrifugation at 700xg, 4 °C for 5 min. Membranes were then resuspended in a cold lysis buffer (10 mM Tris, 150 mM NaCl, 100 μM MgCl_2_, 1 mM EDTA, 100 μM phenylmethyl sulfonyl fluoride (PMSF), 15% glycerol) and homogenized for 5 min on ice. The lysis mixture was then incubated while rotating for 30 min at 4 °C. A second homogenization was carried out, and the lysis mixture was centrifuged at 33,300xg for 25 min, 4 °C. The supernatant was discarded, and pellets were dissolved in solubilization buffer (20 mM Tris, 150 NaCl, 15% glycerol, 2% (w/v) n-dodecyl-β-D-maltoside (DDM; Anatrace) and 0.2% (w/v) Cholesterol Hemisuccinate, Tris Salt (CHS; Anatrace)). The solubilization mixture was then homogenized and incubated for 1 h, after which it was centrifuged at 370,000xg for 25 min, 4 °C. The supernatant of this solubilization step was then incubated for 2h, 4 °C with 60 μl of EZview red anti-Flag beads (Sigma Aldrich). The anti-Flag beads were then spun down (800xg, 2 min) and washed by subsequently decreasing amounts of glycerol to 1% and DDM to 0.03%. After the last washing step, the protein was eluted on ice for 30 min using 100 μl of 0.40 mg/ml of 1× Flag peptide (Genescript) in a buffer containing 1% Glycerol, 0.03% DDM, 20 mM Tris-HCl pH 8.0, 150 mM NaCl. Eluted protein aliquots were flash-frozen in liquid nitrogen. . Analysis of SBAV WT and its mutants’ protein expression was done by transfecting HEK293T cells with 5⍰g of WT/mutant, followed by lysis at 48h post-transfection using Triton lysis buffer (25 mM HEPES, 150 mM NaCl, 1 mM EDTA, 10 % Glycerol (v/v), 1 % triton-x-100 (v/v)). Pull-down of LASV/SBAV RRRR mutants was done by concentrating the protein from the lysate using anti-Flag beads. For western blot analysis, anti-Flag primary antibody (Cell Signalling) was used at a 1:1,000 dilution, followed by horseradish peroxidase (HRP)-conjugated anti-Rabbit secondary antibody (Jackson) at a 1:10,000 dilution. For load control, the membrane was stripped and re-blotted using anti-a-Tubulin DM1A (Millipore) at 1:5000 dilution, followed by horseradish peroxidase (HRP)-conjugated anti-Mouse secondary antibody (Jackson) at a 1:10,000 dilution.

### Cryo-EM image acquisition and 3D reconstruction

A total of 3.5 μl of purified SBAV spike samples (WT or H157M) were applied on glow-discharged (8 s, 12 mA; Pelco easiGlow, Ted Pella) graphene oxide Quantifoil copper grids, R1.2/1.3, (Electron Microscopy Sciences) using a Vitrobot system (Thermo Fischer/FEI) (3.0 s blotting time, 4 °C, 100% humidity). Samples were incubated on the grid for 1 min before blotting was carried out. For analyzing samples in the presence of potassium, the protein samples were supplemented, prior to applying the samples on grids, with a 100 mM potassium chloride solution at a 1:10 ratio to yield a final concentration of 10 mM potassium. Cryo-EM data were collected on the Titan Krios microscope (FEI) operated at 300 kV, using a Gatan K3 direct detection camera. The beam size was 705 nm diameter (fringeless illumination), the exposure rate was 18 e s^−1^ pixel^−1^, and movies were then obtained at 105,000× magnification with a pixel size of 0.824 Å. The nominal defocus range was −0.8 to −2.0 μm. Data processing was carried out with the cryoSPARC v4.1.2 suite ^50^. Patch motion correction and patch CTF estimation were carried out using cryoSPARC Live. Particles were extracted using a 256-pixel box size, and the data sets were cleaned and classified as illustrated for each structure (Extended Data Figs. 1, 7, 13, 14). Final maps were obtained from non-uniform refinement, imposing C3 symmetry.

### Model building, refinement, and analysis

The initial model was generated using ModelAngelo ^51^. This initial model was then manually completed and refined using Coot ^52^ and real-space refinement in Phenix ^53^. Structural analysis and representation were done using PyMol ^54^, ChimeraX ^55^, and CCP4 ^56^.

### Pseudoviral Particle Production and Infectivity Assays

MLV virus-like particles (VLPs) pseudotyped expressing SBAV GPC were produced by transfecting retroviral transfer vector pLXIN-Luc encoding luciferase as a reporter gene together with SBAV-flag or mutated SBAV-flag in pcDNA3.1 into the GP2-293 retroviral packaging cell line (Clontech). GP2-293 cells were seeded at 5×10^6^ on 10-cm plates and transfected 24 h later with 5 μg of SBAV-Flag or SBAV-Flag-mut and 5 μg Luciferase using Lipofectamine 2000 (Invitrogen). Cells’ media were replaced 5 h later to full medium, i.e., DMEM (Biological Industries) supplemented with 1% Pen-Strep (v/v), 1% Glutamine (v/v), and 1% sodium pyruvate (v/v). At 48 h post-transfection, media containing pseudoviruses were harvested, and VLPs were concentrated 10 times by the addition of PBS PEG 6000 8 % (w/v) (Sigma), followed by incubation at 4º C for 24h, centrifugation at 10,000xg for 20 min and resuspension in full medium. The concentrated VLPs were stored at - 80º C until use.

For infectivity assays, HEK293T cells stably expressing human TfR1 ^18^ were seeded on a poly-L-lysine-pre-coated white, chimney 96-well plate (Greiner Bio-One). Cells were left to adhere for 3 h, followed by the addition of SBAV VLPs. Cells were washed from the viruses at 18 h post-infection, and luminescence from the activity of luciferase was measured at 48 h post-infection using a Tecan Infinite M200 Pro plate reader (TECAN) after applying Bright-Glo reagent (Promega) to the cells.

To quantify the relative levels of VLPs, we used RT-qPCR to determine the RNA levels of the Luciferase reporter gene. VLPs containing Media were centrifuged at 13,000 rpm 4° C for 10 min, and 50 ⍰l from the supernatants were treated with 2 ⍰g/ml RNAse A (Bio Basic Inc) for 10 min at RT. To inhibit RNAse A activity, 40 units of RNasin® Ribonuclease inhibitor (Promega) were added, and the reaction was incubated for 10 min at 37° C. RNA was then extracted from viral particles using RNAeasy mini kit (Qiagen). To eliminate remnants of the Luciferase-containing retroviral vectors, RNA was treated with RNAse-free DNAse I (NEB) for 10 min at 37 °C, followed by heat inactivation at 75 °C for 10 min. Thereafter, cDNA was prepared from RNA using a High capacity cDNA reverse transcription kit (Applied Biosystems). cDNA was diluted 1:40 and subjected to quantitative PCR (qPCR) using specific primers for GFP and Fast SYBR green master mix (Applied Biosystems).

### FACS analysis

HEK293T cells were seeded at 5×10^6^ on 10-cm plates and transfected 24 h later with 5 μg of LASV-flag/SBAV-flag/SBAV-flag-H157M/H300A using Lipofectamine 2000. Cells’ media were replaced 5 h later to full medium. At 48 h post-transfection, cells were washed and scraped in PBS and placed on ice. The cells were washed by centrifugation at 600x g for 1 min and re-suspended in PBS supplemented with 3 % BSA, followed by 2 wash steps in PBS and incubation for 30 min with 10 mg/ml of Arenacept (sAD Neotomis TfR1 Fc) diluted in PBS with 1 % BSA. Cells were then washed and incubated with a 1:500 dilution of goat anti-human Cy3 conjugated secondary antibody (Jackson) for 15 min. LASV-transfected cells (and un-transfected cells) were used as a negative control. Before running the FACS analysis, cells were washed twice in PBS and filtered through a 35 μm cell strainer (Corning). FACS analysis was performed using a ZE5 flow cytometer (BIO-RAD) and FlowJo cell analysis software (FlowJo, LLC, Ashland, Ore.).

## Acknowledgments

The Diskin lab is supported by research grants from the Ernst I. Ascher Foundation, Ben B. and Joyce E. Eisenberg Foundation, Estate of Emile Mimran, Jeanne and Joseph Nissim Center for Life Sciences Research, Donald Rivin, Stanley and Tanya Rossby Endowment Fund, Dr. Barry Sherman Institute for Medicinal Chemistry, as well as from the Israel Science Foundation (grant No. 209/20).

## Author contributions

R.D. conceived and oversaw this research. H.C-D. produced and purified proteins. H.C-D. & R.D. performed EM analyses and solved structures. H.C-D. performed infectivity assays. H.C-D. & M.K. performed WB analyses. All authors contributed to the data analysis. R.D. & H.C-D. prepared the manuscript.

## Data availability

Coordinates file and experimental density maps were deposited at the PDB and EMDB under accession codes 9FYA/EMD-50862, 9FYE/EMD-50864, and 9FYG/EMD-50865 for the WT SBAV spike in the closed and open conformation and for the SBAV H175M mutant, respectively.

