## Supplementary Information for "Potassium induces conformational changes in the Sabiá virus spike complex"

This file contains:
Extended Figures 1-15
Extended Table 1

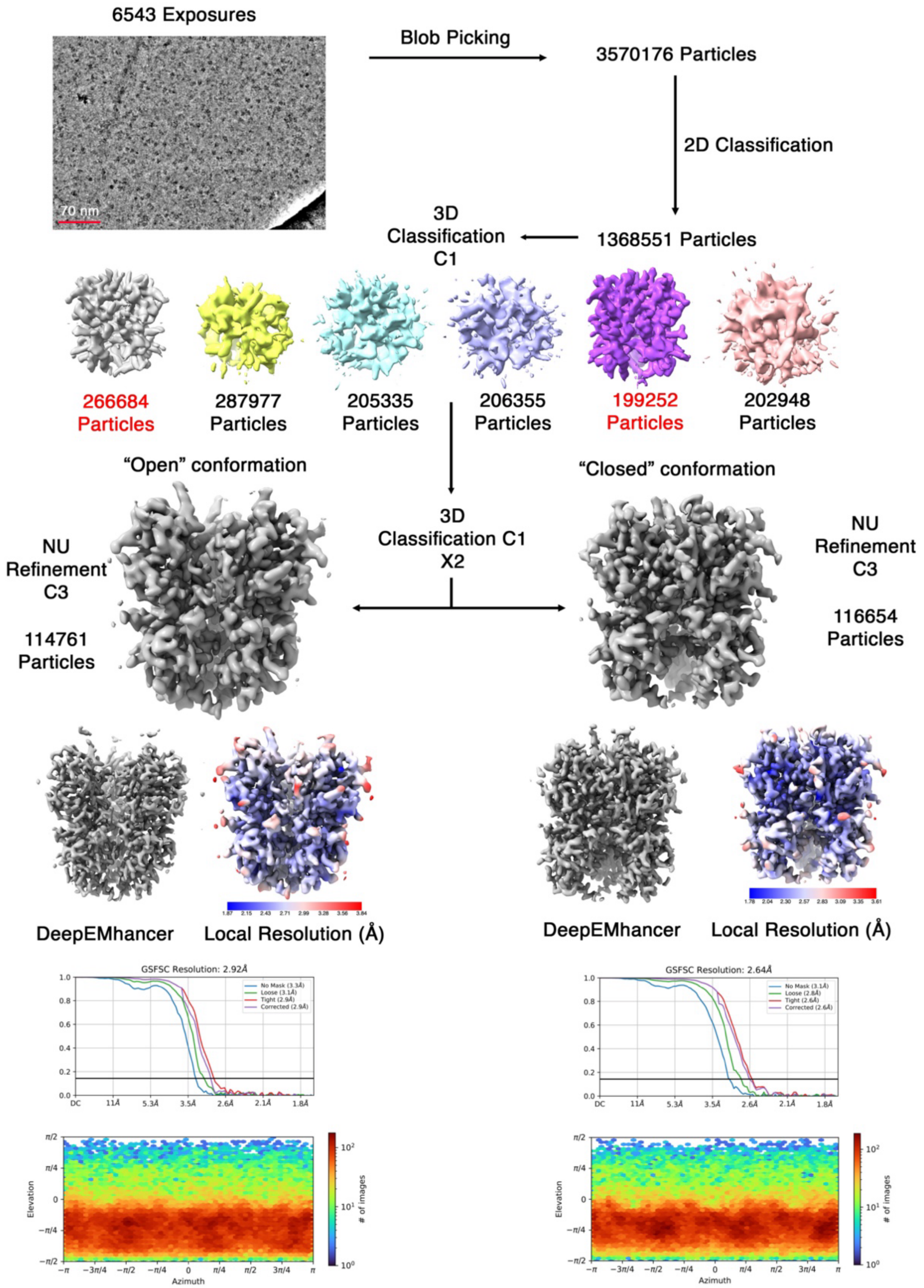


**Extended Data Fig. 1** | Single particle EM reconstruction workflow. Particles noted in red were carried over for subsequent steps.

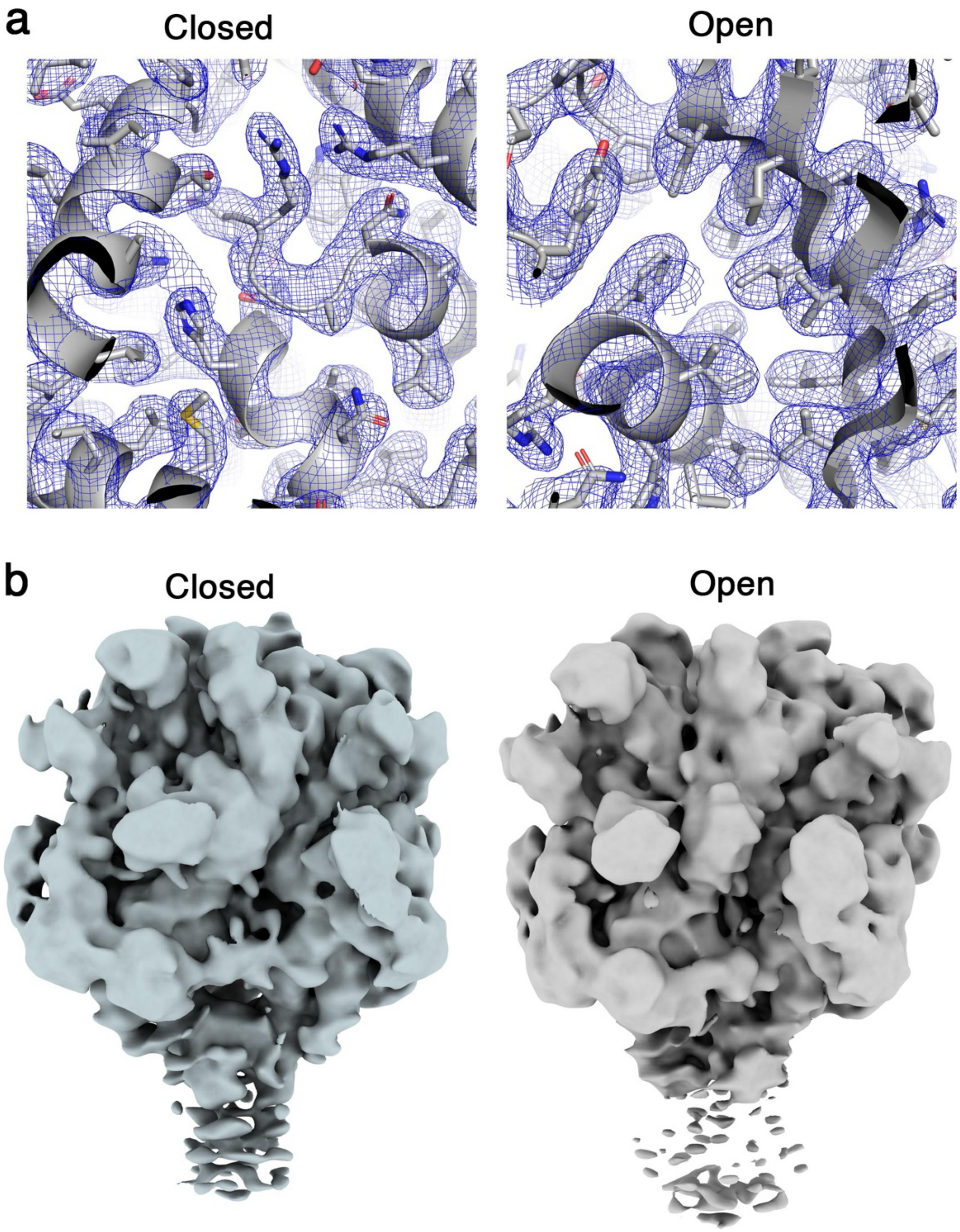


**Extended Data Fig. 2** | Quality of density. **a**. Representative sections of the EM maps in the open and closed states. Maps are shown as blue meshes at $\sigma$=5.0. **b**. Density for the TM region in the closed and open conformations. Low-pass filtered maps at 6 Å resolution are shown at a contour level of 0.02. Some density for the TM region is visible in the closed configuration but is not sufficiently detailed for model building.

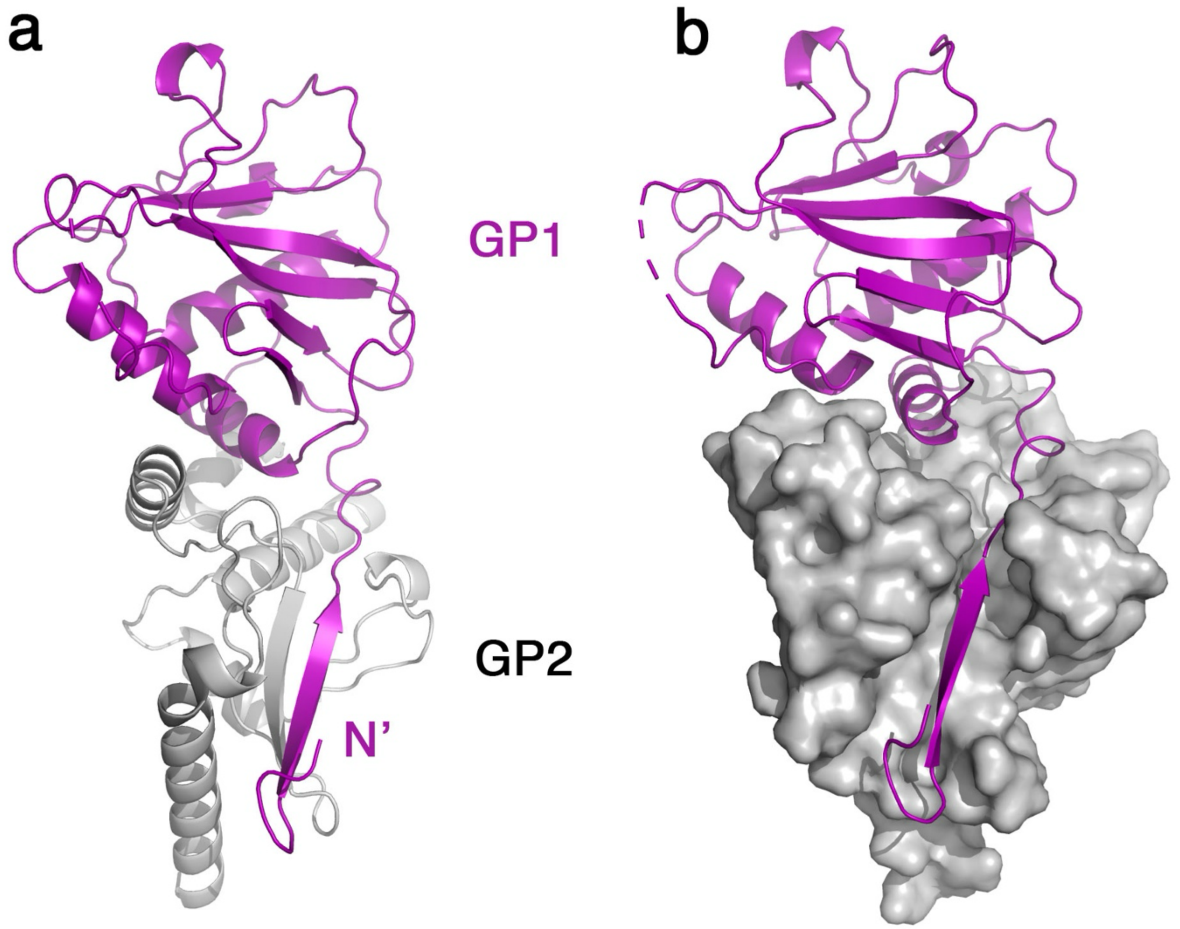


**Extended Data Fig. 3** | The N-terminus of GP1. **a**. Ribbon diagrams of GP1 (purple) and GP2 (grey) are shown, and the N-terminus of GP1 is indicated. **b**. GP2 is shown using a surface representation, illustrating the pocket on GP2 that accommodates the N-terminus of GP1.

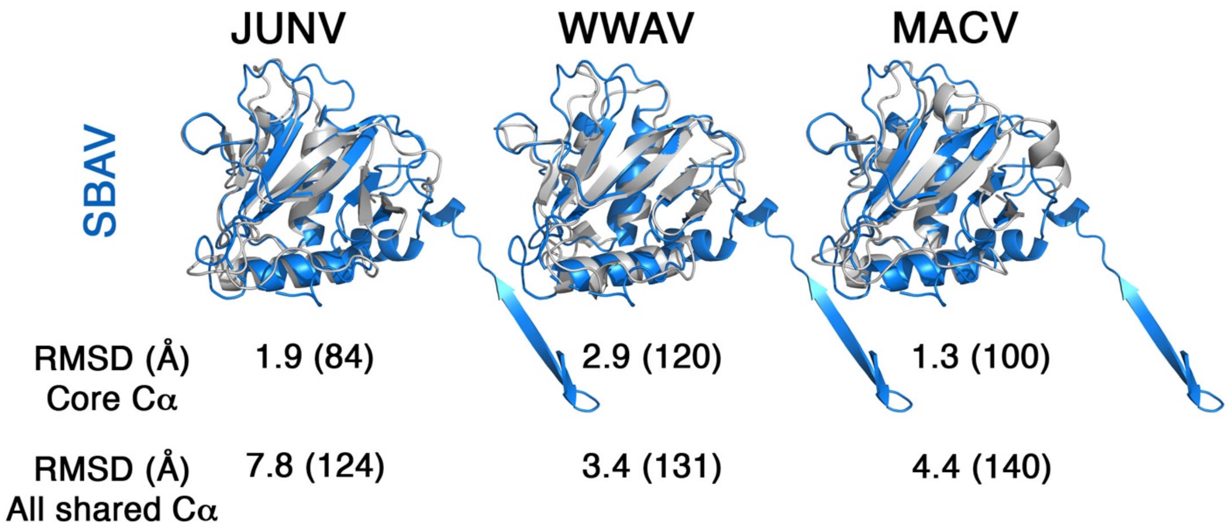


**Extended Data Fig. 4** | Comparison of the SBAV GP1 domain with GP1 domains of other NW arenaviruses. Superimposition of GP1 from SBAV (blue) with the GP1 domains of JUNV (PDB: 5EN2), WWAV (PDB: 5NSJ), and MACV (PDB: 6S9J). RMSD values, calculated for the conserved cores or for all Cα atoms (numbers of Cα atoms are indicated in parentheses) are shown.

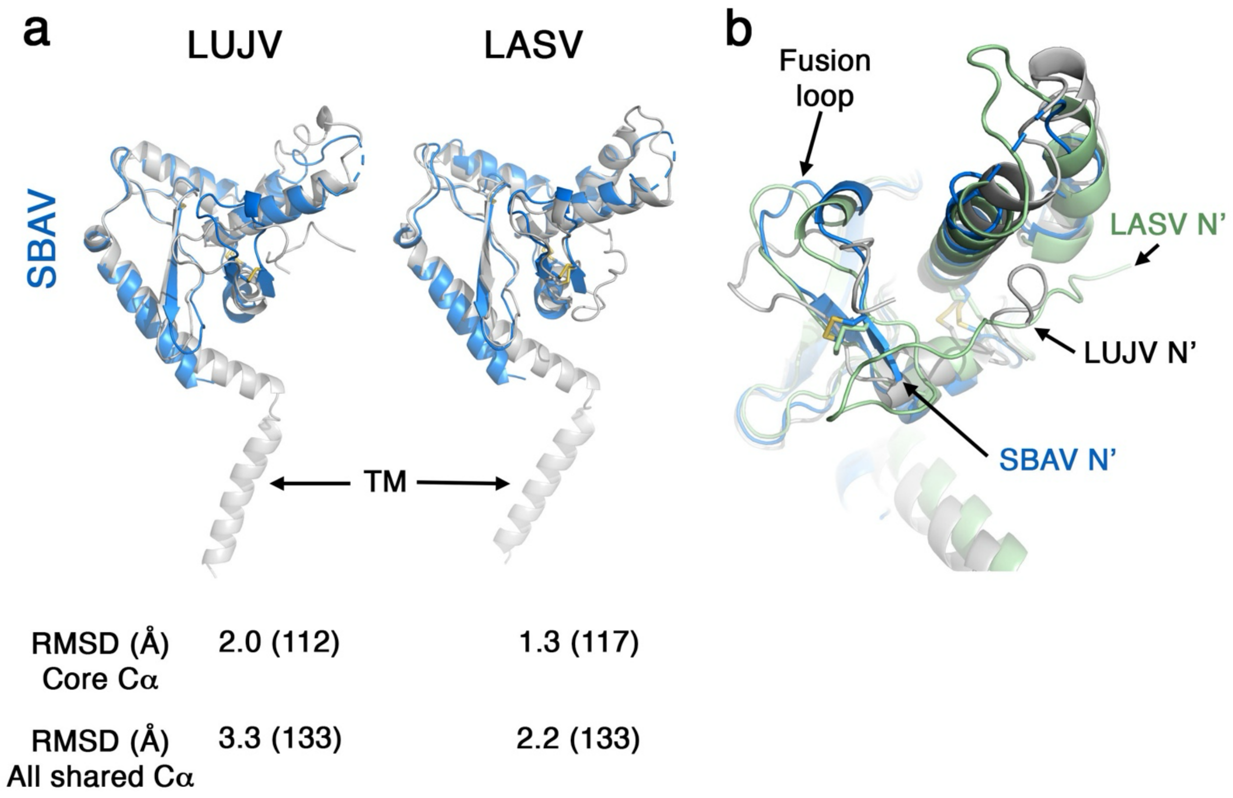


**Extended Data Fig. 5** | Comparison of GP2 domains. **a**. Superimposition of the SBAV GP2 with the GP2 domains of LASV (PDB: 7PUY) and JUNV (PDB: 8P4T). The RMSD values for all Cα atoms or a subset of Cα atoms, as indicated in parentheses, are shown. **b**. A focused view of the fusion loop and the extent of the N-termini regions that were modeled in SBAV, LASV, and LUJV. Disulfide bonds are shown as sticks.

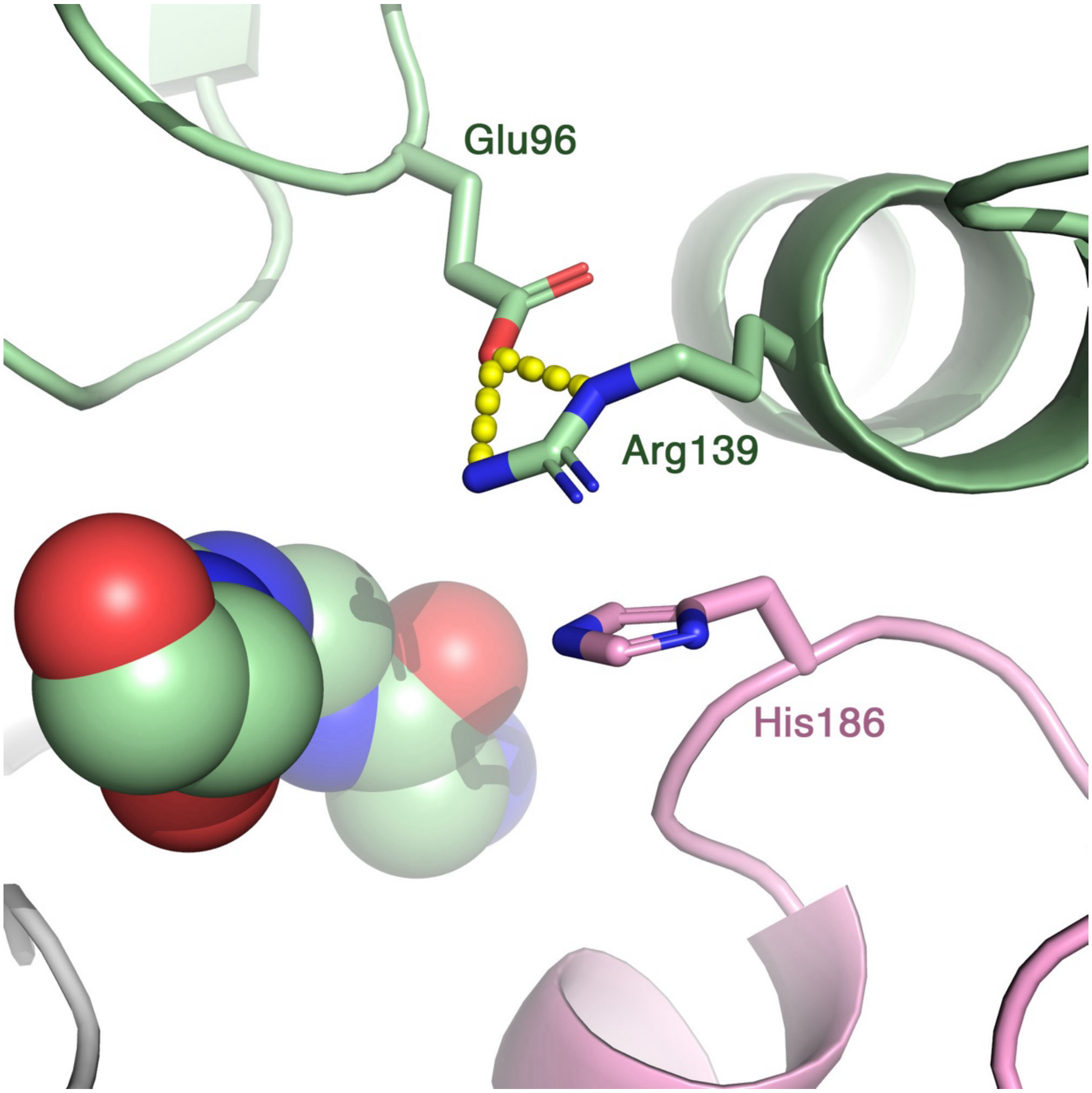


**Extended Data Fig. 6** | The interaction between His186 and Arg139 in the closed state. The interface between two GP1 domains (pink and green) is shown. Arg139 is positioned by forming a salt bridge with a nearby Glu96 on the same GP1 monomer. The imidazole ring of His186 is almost parallel to the guanidino group of Arg139, forming $\pi$-$\pi$ stacking or $\pi$-cation interaction. The aliphatic portions of both residues form van Der Waals interactions.

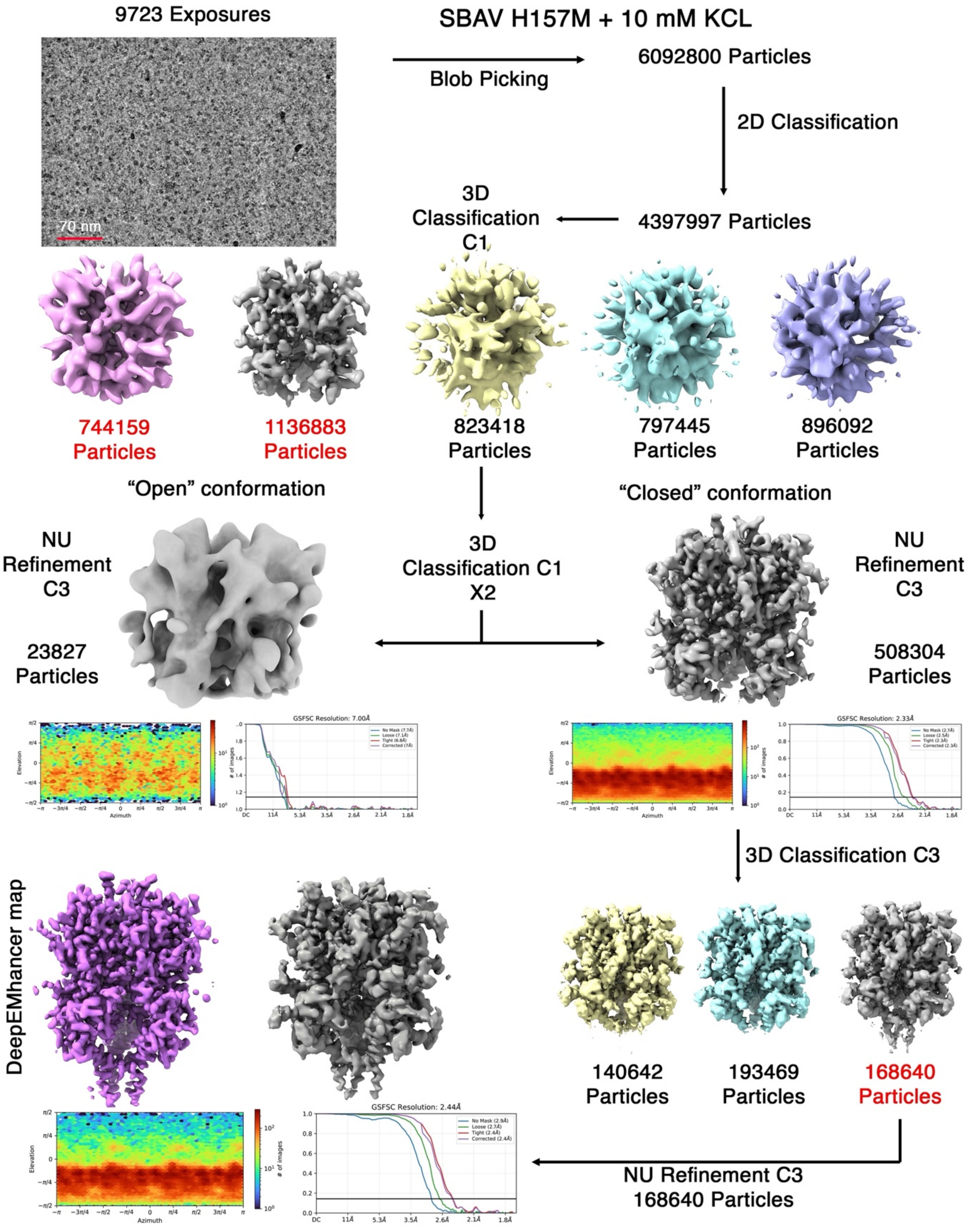


**Extended Data Fig. 7** | Single particle EM reconstruction workflow for SBAV H157M in the presence of 10 mM potassium. Particles noted in red were carried over for subsequent steps.

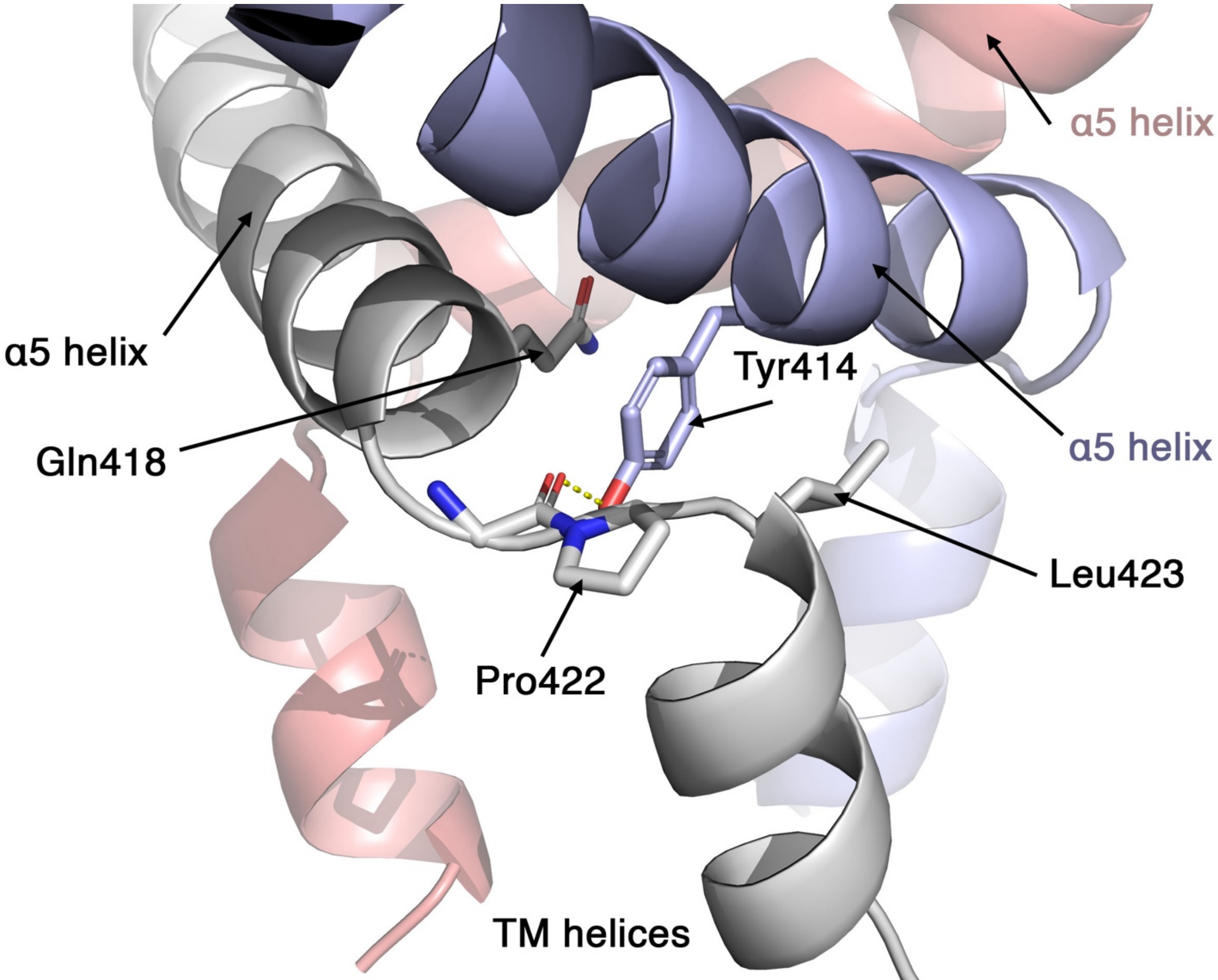


**Extended Data Fig. 8** | The interaction between the α5 helices in the closed state. Each GP2 chain is represented in a different color. A hydrogen bond is shown with a dashed yellow line. The transmembrane (TM) helices are noted.

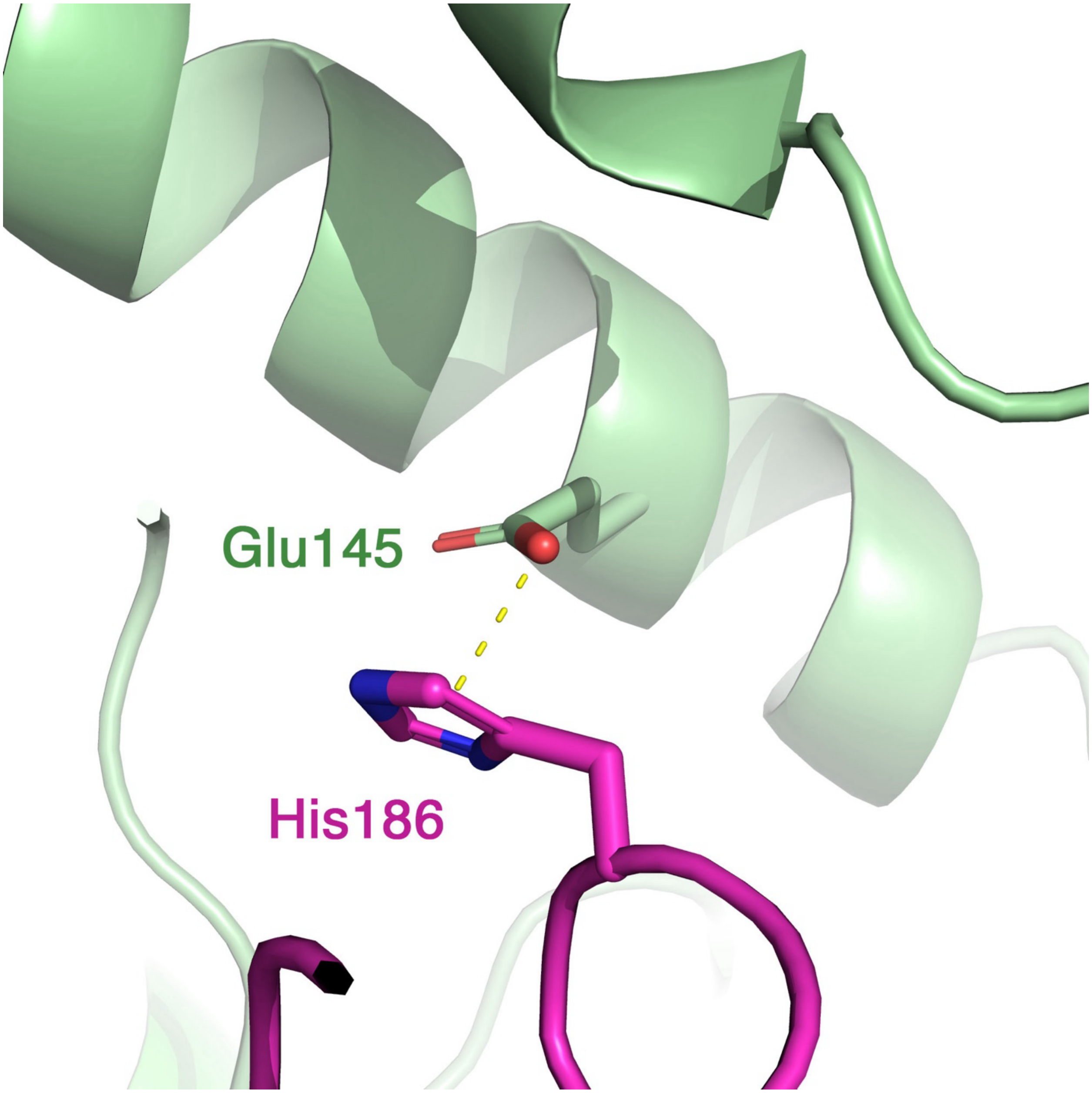


**Extended Data Fig. 9** | Interaction between His186 and Glu145. An oxygen atom of the Glu145 carboxyl group is located 3 Å from the center of the imidazole ring of His186.

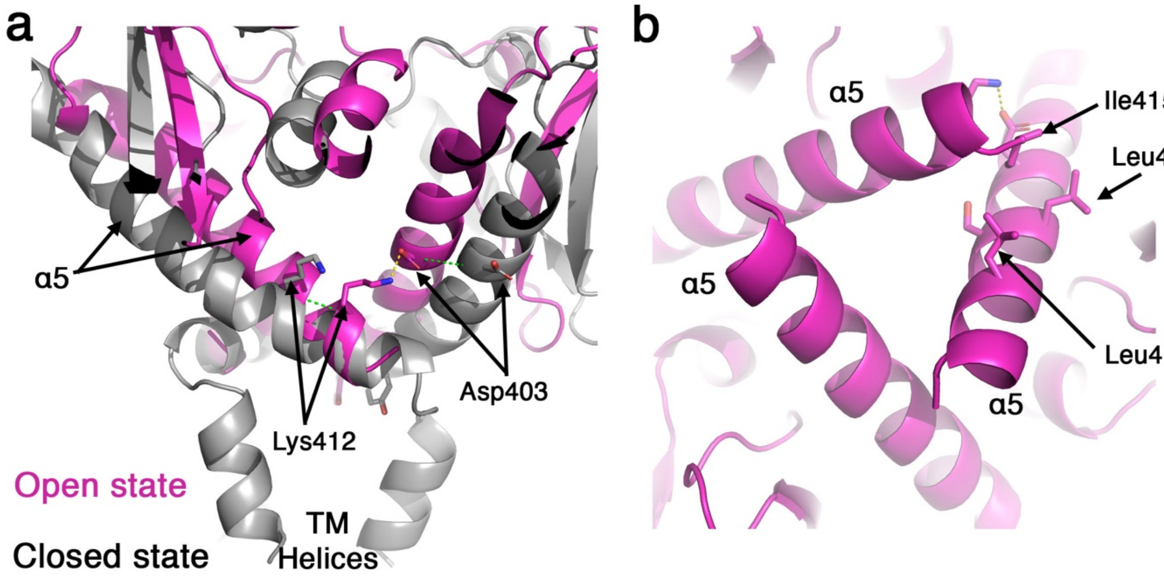


**Extended Data Fig. 10** | The transition of the $\alpha$5 helices between the closed and open states. **a**. The closed (grey) and open (purple) states are shown from a 'side' view. The movement of the C$\alpha$ atoms of Lys412 (6 Å) and Asp403 (8 Å) between the states is illustrated with green dashed lines. Salt-bridge interaction is shown with a yellow dashed line. **b**. A 'bottom' view of the $\alpha$5 helices in the open state. The interaction between the $\alpha$5 helices is shown.

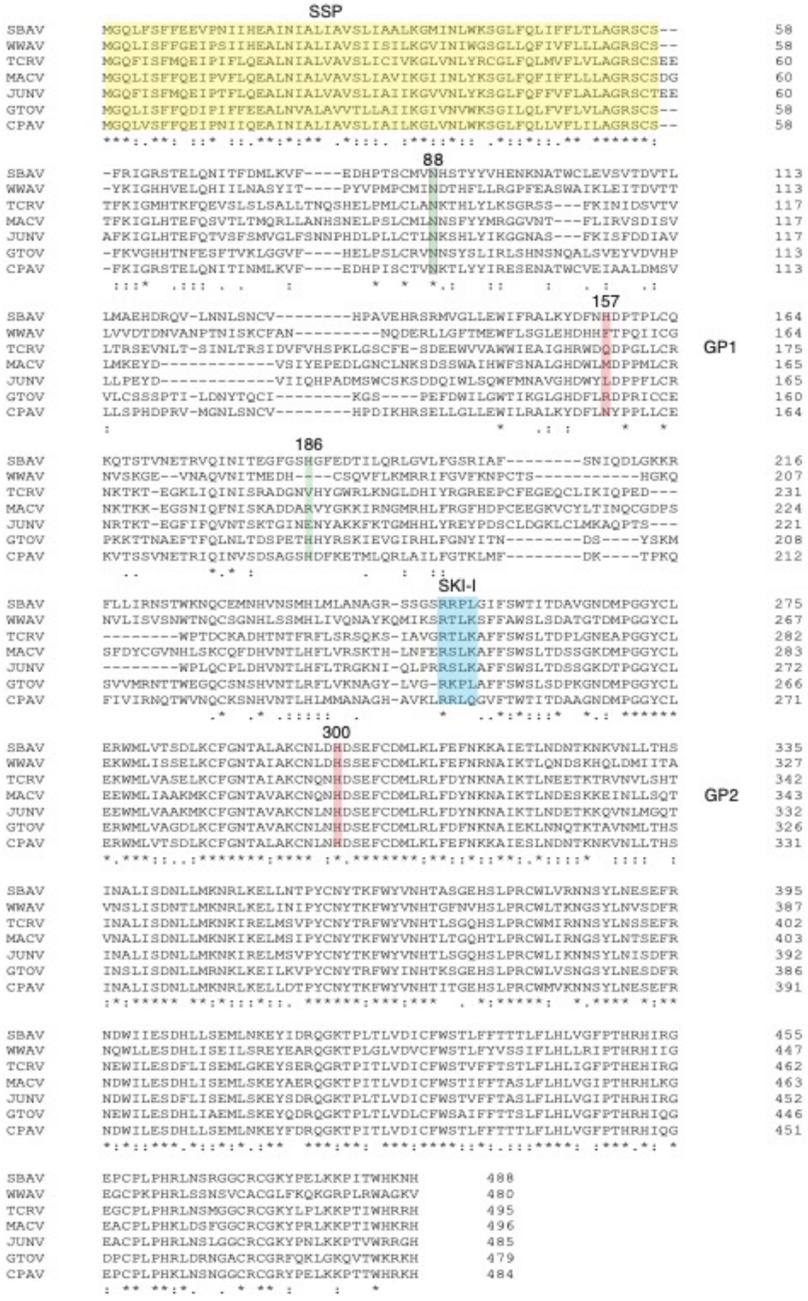

**Extended Data Fig. 11** | Multiple sequence alignment of the GPCs from the NW Sabiá virus (SBAV), Junín virus (JUNV), Machupo virus (MACV), Guanarito virus (GTOV), Chapare virus (CHAPV), Whitewater Arroyo virus (WWAV), and Tacaribe virus (TCRV) (Accession codes: YP_089665.1, P26313.2, NP_899212.1, NP_899210.1, YP_001816782.1, YP_001911113.1, and NP_694849.1, respectively). The SSP, GP1, and GP2 are indicated, as well as the SKI-I site and the residues of interest.

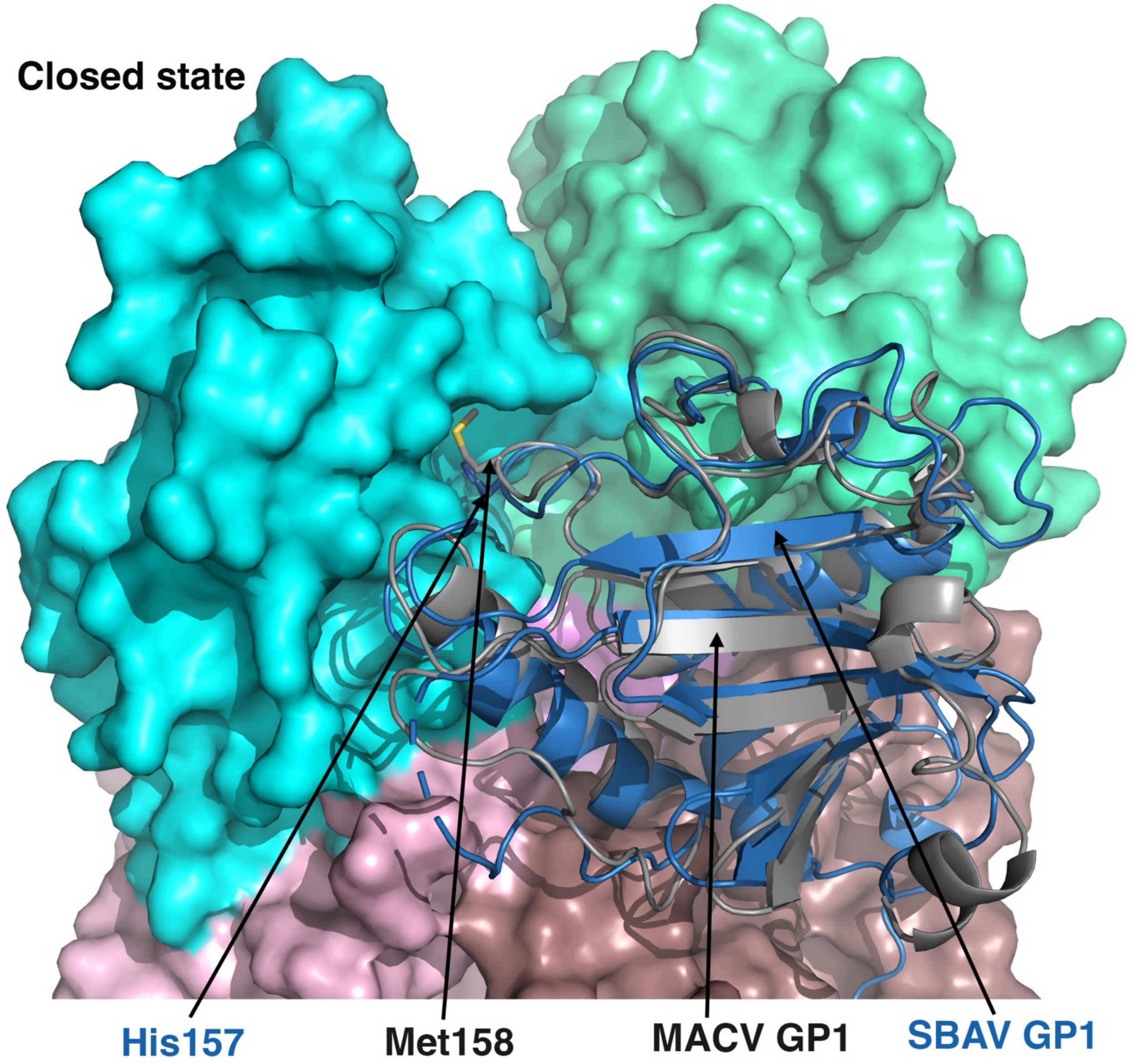


**Extended Data Fig. 12** | Methionine in position 157 can substitute His157 and fit into the hydrophobic pocket. MACV GP1 (PDB: 3KAS) is superimposed on one SBAV GP1 (blue) in the closed state of the spike. Surface representations of the neighboring GP1 domains show the cavity that accommodates His157.

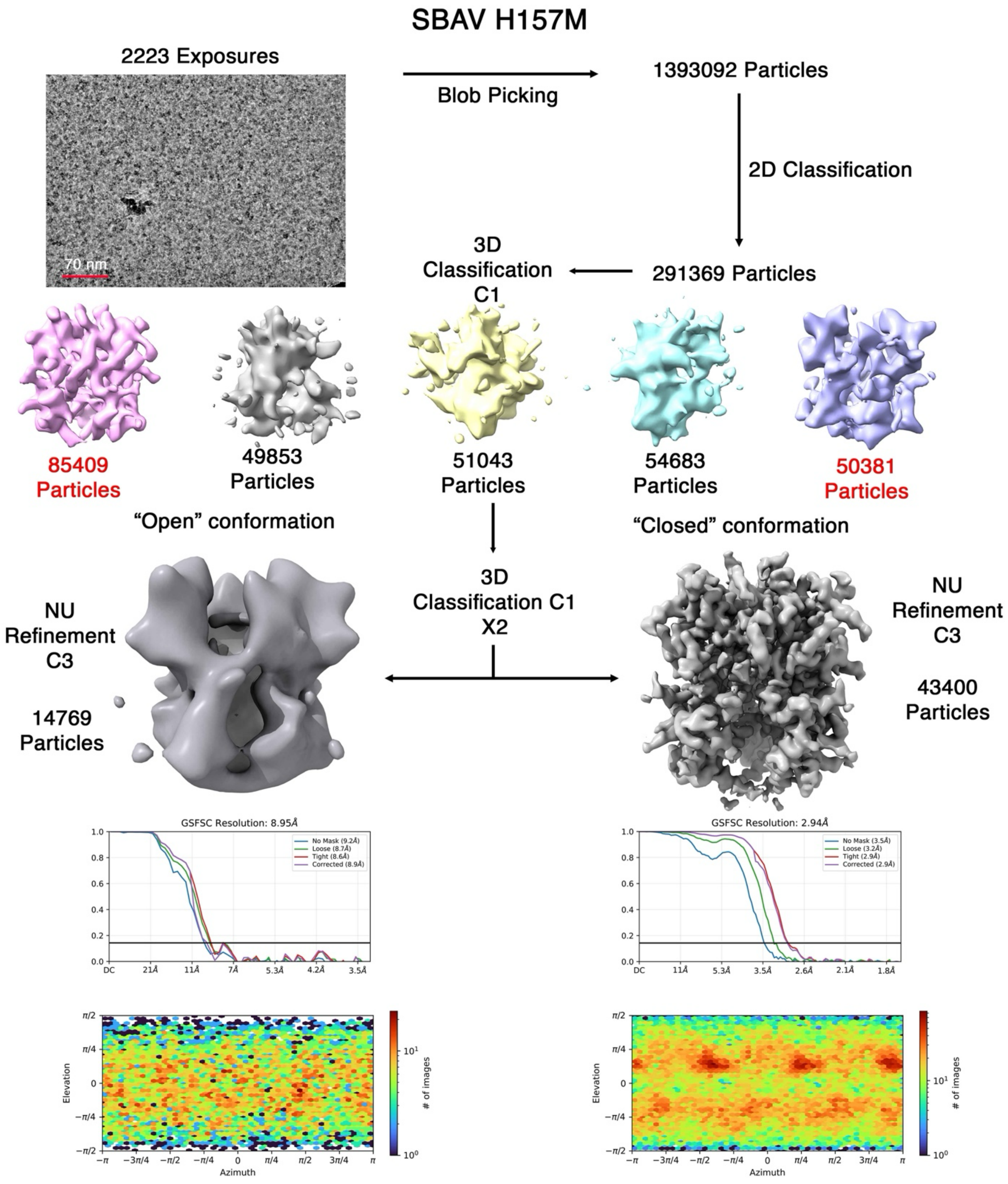


**Extended Data Fig. 13** | Single particle EM reconstruction workflow showing the reconstruction of the SBAV H157M mutant. Particles noted in red were carried over for subsequent steps.

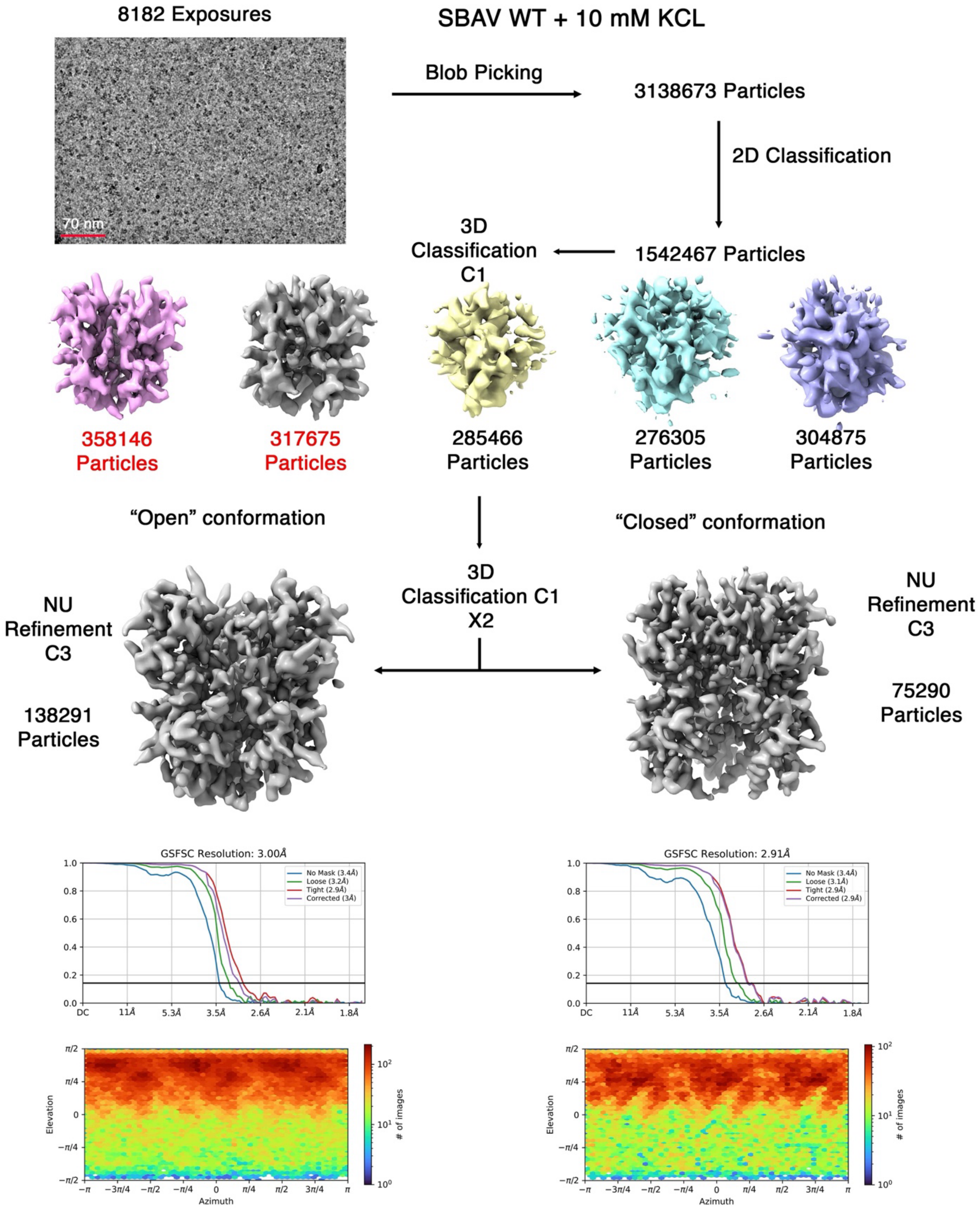


**Extended Data Fig. 14** | Single particle EM reconstruction workflow of the SBAV spike in the presence of 10 mM potassium. Particles noted in red were carried over for subsequent steps.

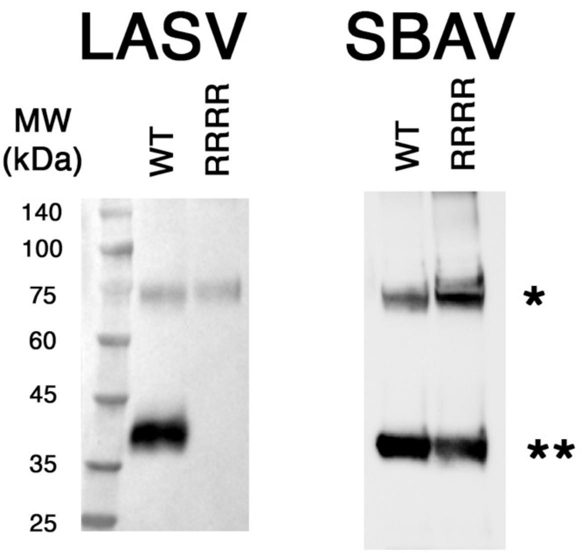


**Extended Data Fig. 15** | WB analysis of SBAV and LASV GPCs with substitutions of their SKI-I recognition sites with an "RRRR" furin recognition sequence. Samples were concentrated with anti-Flag beads, and the GPCs were detected using an anti-Flag antibody. The location of the GPC precursors and the cleaved GP2s are indicated with '*' and '**', respectively.

| | WT | | H157M |
|---|---|---|---|
| | Closed (9FYA, EMD-50862) | Open (9FYE, EMD-50864) | Closed (9FYG, EMD-50865) |
| Model | | | |
| Composition(#) | | | |
| Chains | 11 | 11 | 14 |
| Atoms | 8893 (Hydrogens: 0) | 9100 (Hydrogens: 0) Protein: | 10147 (Hydrogens: 0) |
| Residues | Protein: 963 Nucleotide :0 | 963 Nucleotide: 0 | Protein: 1020 Nucleotide: 0 |
| Water | 3 | 3 | 756 |
| Ligands | ZN: 1<br>NAG: 54<br>BMA: 15<br>MAN: 12 | K: 1<br>BMA: 18<br>NAG: 54<br>MAN: 24 | ZN: 1<br>NAG: 54<br>BMA: 15<br>MAN: 12 |
| Bonds (RMSD) | | | |
| Length (Å) (# > 4σ) | 0.005(0) | 0.004(0) | 0.003(0) |
| Angles (°) (# > 4σ) | 0.805(3) | 0.702(0) | 0.602(0) |
| MolProbity score | 1.42 | 1.5 | 1.36 |
| Clash score | 7.76 | 9.45 | 6.52 |
| Ramachandran plot(%) | | | |
| Outliers | 0 | 0 | 0 |
| Allowed | 0.32 | 0.85 | 1.51 |
| Favored | 99.68 | 99.15 | 98.49 |
| Rama-Z (Ramachandran plot Z-s core,RMSD) | | | |
| whole (N = 939) | 0.99(0.27) | 0.84(0.24) | 0.67(0.26) |
| helix (N = 315) | 0.70(0.31) | 1.40(0.25) | 0.40(0.26) |
| sheet (N = 102) | 2.39(0.50) | 0.91(0.51) | 1.00(0.50) |
| loop (N = 522) | 0.40(0.27) | -0.04 (0.24) | 0.54(0.28) |
| Rotamer outliers(%) | 0.79 | NA | 0.11 |
| Cβ outliers(%) | NA | | NA |
| Peptide plane(%) | | | |
| Cis proline/general | 0.0/0.0 | 0.0/0.0 | 0.0/0.0 |
| Twisted proline/general | 0.0/0.0 | 0.0/0.0 | 0.0/0.0 |
| CaBLAM outliers(%) | 0.66 | 0.44 | 0.51 |
| ADP (B-factors) | | | |
| Iso/Aniso(#) | 8893/0 | 9100/0 | 10147/0 |
| min/max/mean | | | |
| Protein | 0.82/111.73/28.59 | 4.70/115.54/46.59 | 0.34/158.84/38.76 |
| Nucleotide | --- | --- | --- |
| Ligand | 0.96/112.13/40.98 | 5.32/151.66/77.52 | 10.40/133.61/54.84 |
| Water | 67.93/84.53/75.67 | 43.30/59.00/52.18 | 5.23/86.99/34.68 |
| Occupancy | | | |
| Mean | 1 | 1 | 1 |
| occ = 1(%) | 100 | 99.47 | 99.53 |
| > 0occ < 1(%) | 0 | 0.53 | 0.47 |
| occ > 1(%) | 0 | 0 | 0 |
| Data | | | |
| Box | | | |
| Lengths (Å) | 100.53 ,93.11 ,88.17 | 103.00 ,98.88 ,97.23 | 121.13 ,92.29 ,88.99 |
| Angles(°) | 90.00 ,90.00 ,90.00 | 90.00 ,90.00 ,90.00 | 90.00 ,90.00 ,90.00 |
| Supplied Resolution (Å) | 2.6 | 2.9 | 2.4 |
| Resolution Estimates (Å) | Masked Unmasked | Masked Unmasked | Masked Unmasked |
| d FSC (half maps; 0.143) | --- --- | --- --- | --- --- |
| d 99 (full/half1/half2) | ---/---/2.7 ---/---/2.7 | ---/---/3.0 ---/---/3.0 | ---/---/2.6 ---/---/2.6 |
| d model | 2.7 2.7 | 3 3 | 2.5 2.5 |
| d FSC model(0/0.143/0.5) | 2.5/2.6/2.7 2.5/2.6/2.7 | 2.6/2.7/2.9 2.6/2.7/2.9 | 2.0/2.4/2.5 2.0/2.4/2.5 |
| Map min/max/mean | -1.69/2.21/0.02 | -1.56/2.88/0/02 | -1.95/3.04/0.02 |
| Model vs. Data | | | |
| CC (mask) | 0.87 | 0.85 | 0.85 |
| CC (box) | 0.72 | 0.67 | 0.71 |
| CC (peaks) | 0.72 | 0.69 | 0.75 |
| CC (volume) | 0.81 | 0.8 | 0.82 |
| Mean CC for ligands | 0.65 | 0.66 | 0.63 |

**Extended Data Table 1** | Model refinement statistics.
